# Representational similarity supports rapid context retrieval during fear discrimination in ventral hippocampus

**DOI:** 10.1101/2023.09.08.556889

**Authors:** Robert R. Rozeske, Léonie Runtz, Alexandra T. Keinath, Aaron Sossin, Mark P. Brandon

## Abstract

The dorsal and ventral regions of the hippocampus (dCA1 and vCA1) are critical for contextual fear conditioning tasks, yet the dynamics of hippocampal neuronal representations during memory acquisition, retrieval, and extinction processes have yet to be fully elucidated. The canonical theory is that the hippocampus generates and retrieves spatial maps of contexts during memory acquisition and retrieval. It is hypothesized the hippocampus prevents memory interference by generating context representations that are dissimilar and the prediction follows that representation dissimilarity facilitates discrimination between contexts. Here, we developed a context fear memory retrieval task and combined it with 1-photon neuronal imaging in dCA1 and vCA1 of freely behaving mice to test this prediction. We identified several phenomena specific to vCA1. First, fear conditioning induced an immediate and strong representational change that was predictive of freezing behavior. During context discrimination, vCA1 representations of the threatening and neutral contexts became more similar. Third, during threatening context memory retrieval, vCA1 expressed rapid and strong context representations. These unexpected results suggest that representational similarity in vCA1 facilitates faster and more efficient network state transitions. In further support of this view, both phenomena of representational similarity and rapid context recall were no longer observed in vCA1 once fear behavior was extinguished. Together, these results reveal that vCA1 unexpectedly generates similar population codes, promoting faster transitions between network states essential for contextual fear memory retrieval.

**Highlights:**

- We established a novel context teleportation task that allows population level hippocampal recordings during key moments of threat memory acquisition, retrieval, and extinction.
- The ventral region of CA1 (vCA1) exhibited the greatest change in neural representation during fear memory acquisition, compared to dorsal CA1 (dCA1).
- After fear conditioning, representations for threatening and neutral contexts were more similar in vCA1 compared to dCA1, yet representational similarity in vCA1 supported rapid context memory retrieval.
- Context fear memory extinction reversed the context representation in vCA1 to patterns observed prior to fear conditioning.

**Graphical Abstract:** 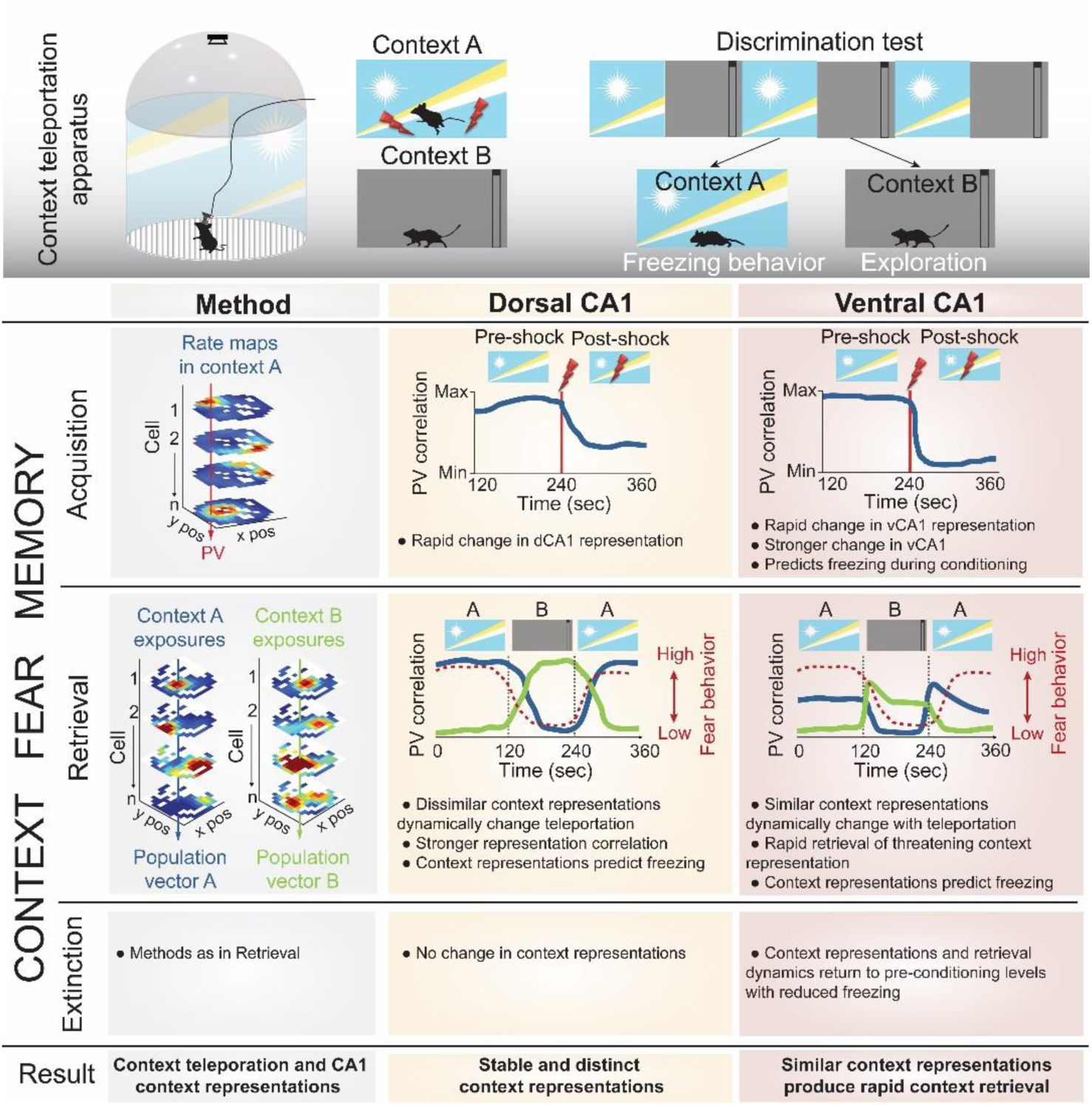

## Introduction

Episodic memory formation in the hippocampus is thought to require the association of a physical context or space ^1,2^ with concurrent non-spatial sensory stimuli ^3–9^. The processes of memory acquisition, retrieval, and extinction enable organisms to generate context-informed predictions based on prior experiences to guide appropriate behaviour ^10–12^. The hippocampus has the dual challenge of preventing memory interference by distinguishing similar experiences into separate memory traces ^13,14^ and updating memory traces to incorporate the organism’s most recent experience ^5,9,15–17^. A wealth of studies demonstrate that threatening memories are encoded as engrams across distributed systems in cortical and subcortical structures, including the hippocampus and basolateral amygdala ^18–20^. Crucially, these memory representations, particularly those associated with threat, are not static but undergo modification through experiences such as stress, memory retrieval, and extinction ^21–23^, consequently altering fear and anxiety expression.

Evidence from neuronal recordings in rodents demonstrate that spatially tuned neurons throughout the hippocampus, known as *place cells* ^24^, effectively form a cognitive map of an environment ^25^. To prevent interference across environments, these place cells undergo *remapping,* creating unique coding schemes to represent distinct context memories ^16,17,26,27^. Accordingly, after remapping the hippocampal context representations become more dissimilar ^2,26,28^. The magnitude of representational dissimilarity after remapping reflects spatial distinctions between the contexts, whereas representational similarity reflects generalizations in the spatial code ^29^. Although remapping is a coding scheme that reduces spatial representation similarity, the process is also evident following threatening experiences ^9,30^, indicating that place cells not only encode specific contexts, but also adapt their tuning properties when an event modifies the meaning of a context. Spatial coding properties differ across the longitudinal axis of the hippocampus ^31–33^. Place fields expressed by dorsal hippocampal CA1 (dCA1) neurons are smaller and contain greater spatial information compared to the larger fields expressed by ventral CA1 (vCAl) neurons. Despite these contrasting sparse and distributed spatial coding schemes, at the population level dCA1 and vCA1 each generate a reliable spatial representation of a given context ^29^ and lesion studies confirm that damage to dCA1 and vCA1 results in severe context memory loss ^34–36^. This dichotomy in spatial mapping could provide complementary representations that balance memory specificity in dCA1 and generalization in vCA1 ^29^. In addition to spatial firing characteristics, the longitudinal axis of CA1 also exhibits functional dissociation, with ventral regions broadly associated with emotional processing ^32,37–40^. Several studies indicate vCA1 directly codes for, or is required for, emotional behavior in vCA1 ranging from anxiety ^41–43^, active avoidance ^44,45^, auditory fear conditioning ^46,47^, and context fear conditioning ^48–50^. These findings indicate that vCA1 preferentially supports context-based memories that contain a threatening component and may differentially represent dynamic changes in threat ^37,51^.

Given the well-defined spatial coding and functional properties of vCA1 and dCA1, we focused our examination on how these subregion differences would actually manifest during moments of context memory acquisition, retrieval, and extinction. To explore this, we developed a context fear memory retrieval task in mice to quantify spatial representations, their temporal expression dynamics, and memory strength, along the longitudinal axis of the hippocampus. Inspired by previous studies ^14,52^, we constructed an arena that allowed remote manipulation of visual and auditory sensory elements to effectively *teleport* mice to different contexts. We hypothesized that vCA1 population activity would be strongly altered by threat memory acquisition, and during memory retrieval vCA1 population activity would rapidly represent the threatening context. Our experimental design also allowed us to examine the relationship of context representation similarity and successful context fear discrimination.

We found that fear conditioning led to increased representational similarity of neutral and threatening contexts in vCA1. During context fear memory retrieval, we observed vCA1 rapidly retrieved threatening context representations. Surprisingly, we observed in vCA1, but not dCA1, representational similarity was a positive predictor of successful contextual discrimination. During fear extinction, representational similarity in vCA1 was reduced and returned to levels observed prior to fear conditioning. These results reveal an unpredicted relationship between context representations and memory expression, whereby vCA1 employs a population coding scheme of context representational similarity to bias faster memory retrieval dynamics. These findings underscore the pivotal role of hippocampal CA1 regions in encoding, retrieving, and updating of contextual fear memories.

## Results

### Mice discriminate threatening and neutral contexts during context teleportation

To evaluate the population dynamics in the hippocampus that underlie the encoding, retrieval, and extinction of an aversive memory, we developed a context fear discrimination task that permits instant transitions, or *teleportation*, between contexts in augmented reality. This task was designed to allow us to record the activity of large hippocampal dCA1 and vCA1 neuronal populations during the transitions between a neutral and conditioned context. Using the *teleporter*, sensory elements were manipulated to create different contexts as follows: (1) context A contained light patterned wall, dark flooring, and tonic white noise; (2) context B contained gray patterned wall, light flooring, and tonic 10 kHz pure tone (**Figure 1A-B**).

**Figure 1:**
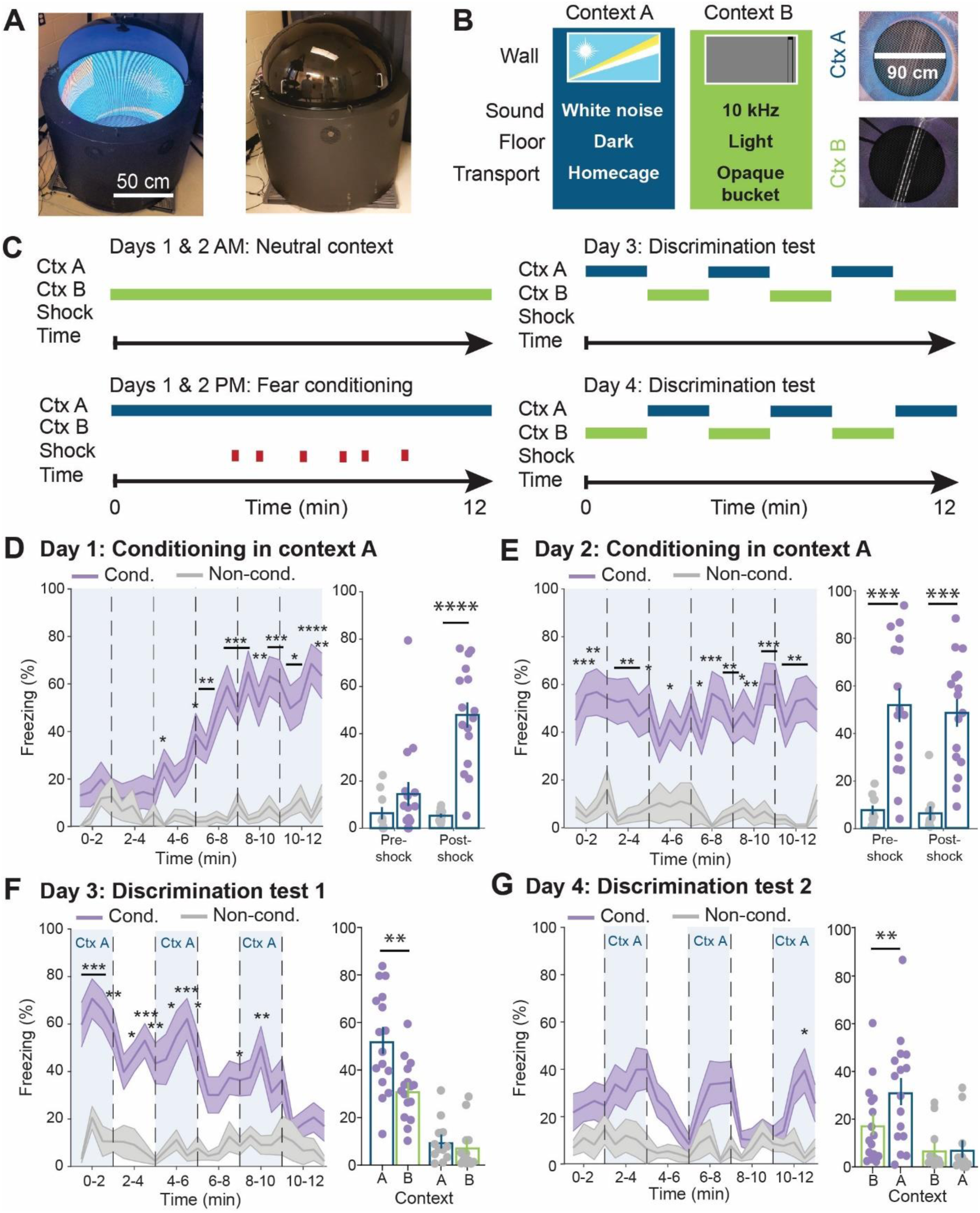
Mice express fear discrimination in novel context retrieval task. **(A)** Aerial view of fear discrimination apparatus with dome removed to display interior (left) and with dome in testing position (right). **(B)** Sensory elements of contexts and mode of transport between colony room and testing suite. **(C)** Experimental protocol: Top and bottom left, on days 1 and 2 mice are exposed to non-shock paired with context B and 6 footshocks paired with context A. Top right, day 3 context discrimination test with alternating context A and B presentations every two minutes. Bottom right, day 4 context discrimination test with alternating context B and A presentations every two minutes **(D-E)** Left, dynamics of freezing behavior in context A on day 1 (**D**) and day 2 (**E**) in conditioned (Cond., n = 16 mice) and non-conditioned (Non-cond., n = 10 mice) groups (day 1: two-way repeated measures ANOVA (RM ANOVA): main effect of group F_(1, 24)_ = 29.29, p < 0.0001; main effect of time F_(7.651, 183.6)_ = 6.433, p < 0.0001; interaction F_(23, 552)_ = 6.034, p < 0.0001 followed by a Bonferroni correction test; day 2: two-way RM ANOVA: F_(1,24)_ = 36.58, p < 0.0001; main effect of time F_(7.81, 187.4)_ = 0.5341, p > 0.05; interaction F_(23, 552)_ = 0.8532, p > 0.05 followed by a Bonferroni correction test). Right, freezing behavior comparison between Non-cond and Cond. groups during the pre-shock period (day 1: Wilcoxon rank-sum: rank = 113.5, p > 0.05; day 2: Wilcoxon rank-sum: rank = 65, p < 0.001) and during the post-shock period (day 1: Wilcoxon rank-sum: rank = 59, p < 0.0001; day 2: Wilcoxon rank-sum: rank = 61, p < 0.001). **(F-G)** Left, dynamics of freezing behavior on day 3 (**F**) and day 4 (**G**) of context fear discrimination task (Cond., n = 16 mice, Non-cond., n = 12 mice). Blue shading represents context A exposure periods (day 3: two-way RM ANOVA: main effect of group F_(1,26)_ = 53.96, p < 0.0001; main effect of time F_(8.937, 232.4)_ = 3.460, p = 0.0005; interaction F_(23, 598)_ = 2.609, p < 0.0001 followed by a Bonferroni correction test; day 4: two-way RM ANOVA: main effect of group F_(1,26)_ = 8.907, p = 0.0061; main effect of time F_(7.660, 199.2)_ = 2.028, p = 0.0476; interaction F_(23, 598)_ = 1.773, p = 0.0148 followed by a Bonferroni correction test). Right, freezing behavior comparison between context A and B exposures in Cond. group (day 3: Wilcoxon signed rank test: rank = 114, p < 0.01; day 4: Wilcoxon signed rank test: rank = 98, p < 0.01) and in Non-Cond. group (day 3: Wilcoxon signed rank test: rank = 46, p > 0.05; day 4: Wilcoxon signed rank test: rank = 8, p > 0.05). All data are expressed as the mean ± standard error of the mean.

During the first two days of the teleporter task, mice were fear conditioned to context A with six footshocks per day, and no footshocks in context B (**Figure 1C, left**). During conditioning, freezing behavior significantly increased in context A, but not context B, in conditioned mice (Cond.), while non-conditioned (Non-cond.) control group expressed minimal immobility (**Figure 1D, Figure S1A-B**). On day 2, high freezing behavior was observed throughout the session in context A, indicating that mice immediately retrieved the fear memory from day 1 (**Figure 1E, Figure S1C-D**).

To examine hippocampal activity during retrieval of the fear memory, on days 3 and 4, we alternated presentations of contexts A and B every two minutes in counterbalanced order (**Figure 1C, right**). In both context teleportation sequences, presentation of context A produced higher freezing levels compared to context B in fear conditioned mice. This context fear discrimination behavior was not observed in non-conditioned mice (**Figure 1F-G, Figure S1E-H**). Furthermore, repeated exposures to context A during discrimination tests generated within-session fear extinction that was examined in subsequent analyses (**Figures 1F-G**, **6A**). Thus, our novel apparatus and behavioral protocol enables the quantification of encoding, retrieval, and extinction of a threatening context memory during rapid context transition.

### Context fear conditioning induces strong remapping of context representation in vCA1

To examine if context fear conditioning alters spatial representations in dCA1 and vCA1, we recorded calcium activity of hippocampal calcium-calmodulin II positive (CaMKII+) neurons during day 1 using microendoscope calcium imaging^53^ (miniscope.org) (**Figure 2A-B, Figure S2A-E**). Given our observation that footshocks significantly change behavior, and thus make traditional rate map comparisons difficult due to lower spatial sampling, we developed an analysis to examine changes to the population-level representation of space across time. To this end, we generated spatial rate maps of each recorded cell using the neuronal activity and locomotor trajectory of the mouse from the period prior to footshock delivery (**Figure 2C, Figure S2I, S2K**). Importantly, we excluded periods of immobility when constructing population vectors to isolate spatial representations of the neuronal population. For each mouse we constructed a matrix that contained the activity of each neuron at each spatial bin (N neurons 𝗑 M spatial bins), termed pre-shock population vector (pre-shock PV)^54,55^. Then, for every *i*^th^ time bin we compared the pre-shock PV for the spatial bin that the mouse occupied during the *i*^th^ time bin, with the PV of neuronal activity during the *i*^th^ time bin, using Pearson’s correlation (see Methods for parameters) (**Figure 2C**) to monitor how the population activity in context A changes throughout the fear conditioning session. Thus, the pre-shock PV correlation during the first four minutes of context A is the *ground truth* hippocampal representation for context A, therefore correlation values in the range of r = 0.3 to 0.4 are considered a perfect match to the pre-shock context representation. We observed in dCA1 and vCA1 that the pre-shock PV of conditioned mice was significantly altered upon footshock, most strikingly in vCA1, while in those same mice it remained stable upon exposure to neutral context B (**Figure 2D, 2F top, Figure S2F**). In contrast, non-conditioned control mice maintained a stable PV in both contexts (**Figure 2D, 2F bottom, Figure S2G-H**). We calculated the within-subject difference in PV correlation between the pre- and post-shock periods to assess the effect of fear conditioning on the pre-shock spatial representation of the context. Indeed, the correlation difference in dCA1 and vCA1 was significantly higher in conditioned mice compared to non-conditioned mice in context A, while remaining stable in context B (**Figure 2E, 2G**). We obtained similar results comparing the spatial rate maps for each neuron during the pre- and post-shock periods (**Figure S2I-L**). The rapid change in the PV correlation following the first shock suggests that a threatening representation is immediately formed by updating spatial representations in both vCA1 and dCA1.

**Figure 2:**
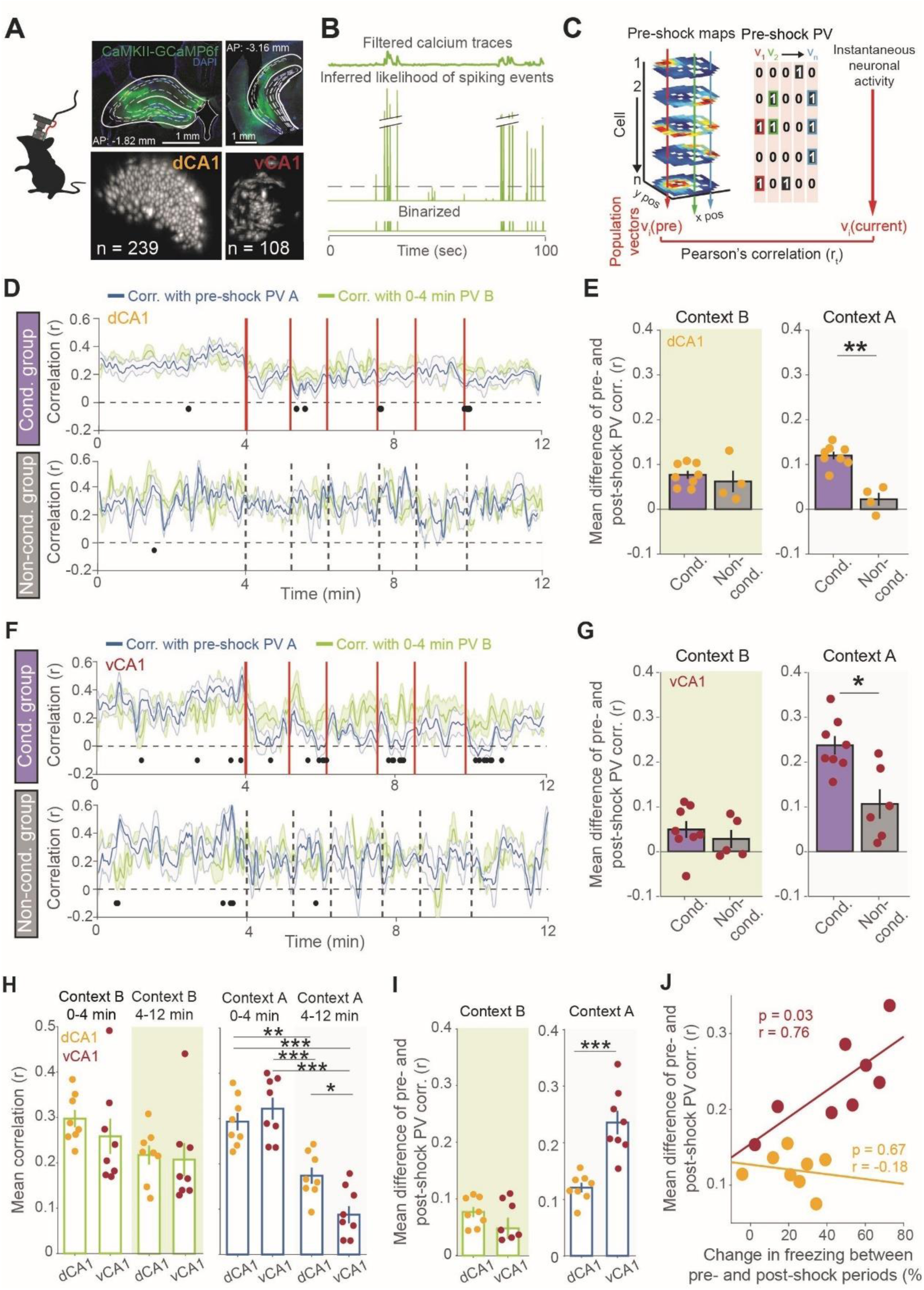
Context fear conditioning produces stronger change in vCA1 representation. **(A)** Freely moving one-photon calcium imaging in dCA1 and vCA1 using a miniscope. Top, coronal brain sections illustrating the GRIN lens location and GCaMP6f expression in dCA1 (left) and vCA1 (right). Bottom, miniscope field of view of the maximum projection of cell segments recorded in dCA1 (left) and vCA1 (right). **(B)** Schematic of filtered neuronal activity extracted from calcium transients (top), the resulting inferred likelihood of spiking events (ILSE, middle) and the signal binarization according the upper 30% of ILSEs (bottom). **(C)** Schematic of the hippocampal representation analysis method. A population vector (PV) was constructed by taking the activity of all recorded cells and organizing it by spatial bin of the fear conditioning apparatus. The PV analysis was performed for the preconditioning period (Pre-shock PV, left). The Pre-shock PV was correlated with the neuronal activity (Pearson’s r) at 2 sec intervals. **(D)** Dynamics of PV correlation from cells recorded in dCA1 during day 1 context A and B exposures in the Conditioning (Cond.) group (top, n = 8 mice) and Non-conditioned (Non-cond.) control group (bottom, n = 4 mice). Vertical red lines indicate shock administration during context A and the corresponding time is represented by dashed lines in the Non-cond. control group. For both groups, no shock was delivered during context B exposure. Black dots indicate timepoints of significant (Wilcoxon signed-rank test, p < 0.05) pairwise comparisons between context A and B correlation values. **(E)** The mean pre-shock PV correlation was calculated for the pre-shock (0-4 min) and post-shock (4-12 min) periods for each mouse, in each group, to assess experience-dependent spatial representation changes. The difference of each mouse’s pre- and post-shock PV correlation was only significant for the Cond. group in context A (context A, Wilcoxon rank-sum: rank = 68, p = 0.004; context B, Wilcoxon rank-rum: rank = 58, p > 0.05). **(F)** Dynamics of PV correlation as in **D** but recorded from vCA1 for Cond. group (top, n = 8 mice) and Non-cond. group (bottom, n = 5 or 6 for context B and A exposures, respectively). **(G)** The difference of the mean pre-shock PV correlations per mouse as in **E**. A significant difference was only observed in the fear conditioned group in context A (context A, Wilcoxon rank-sum: rank = 79, p = 0.013; context B, Wilcoxon rank-rum: rank = 63, p > 0.05). **(H)** Comparison of the mean “pre-shock” PV correlations of mice recorded in dCA1 (n = 8 mice) and vCA1 (n = 8 mice) during context B exposure (left) and context A conditioning (right). Left, no significant differences between vCA1 and dCA1 in non-shocked context B (one-way ANOVA: F_(3,28)_ = 1.705, p > 0.05). Right, in shock-paired context A mean correlations are significantly reduced between pre-shock (0-4 min) and post-shock (0-4 min) periods. Mean correlations are significantly different between vCA1 and dCA1 in context A only during the post-shock period (one-way ANOVA: F_(3,28)_ = 29.07, p < 0.0001 followed by Bonferroni’s multiple comparison test). The statistics presented correspond to Bonferroni’s comparison test. **(I)** Comparison of dCA1 (n = 8 mice) and vCA1 (n = 8 mice) mean correlation differences from 0-4 and 4-12 min is significant in shock-paired context A (right, Wilcoxon rank-sum: rank = 37, p = 0.0003), but not context B (left, Wilcoxon rank-sum: rank = 80, p > 0.05). **(J)** Mean PV correlation difference of pre- and post-shock periods is significantly correlated with the difference in freezing behavior between pre- and post-shock periods only in vCA1 (dCA1: correlation coefficient r = −0.18, p > 0.05, n = 8 mice; vCA1: correlation coefficient r = 0.76, p < 0.05, n = 8 mice). All data are expressed as the mean ± standard error of the mean.

Next, we examined if fear conditioning differentially affects dorsal and ventral hippocampal regions. We compared the pre-shock PV correlation during the pre- and post-shock periods in dCA1 and vCA1. The mean correlation value during the post-shock period (4-12 min) in context A was significantly lower in vCA1 compared to dCA1 (**Figure 2H**). Furthermore, a greater correlation difference between pre- and post-shock periods was observed in vCA1 compared to dCA1 during fear conditioning in context A, and importantly, there was no difference between CA1 subregions during neutral context B exposure (**Figure 2I**). Finally, we observed in vCA1, but not dCA1, that greater alterations in population activity were positively correlated with increased freezing behavior during fear conditioning (**Figure 2J**), an effect that was preserved when cell numbers in vCA1 and dCA1 were matched (**Figure S4A-D**). Together, these results demonstrate that context fear conditioning induces greater change in conditioned context representation in vCA1 and this change is positively correlated with fear memory strength.

### Context representations are more spatially distinct in dCA1 compared to vCA1 during fear discrimination test

To examine the time course of hippocampal spatial representation updating that supports behavioral discrimination of conditioned and neutral contexts, we adopted a similar spatial map PV correlation approach that was used during fear conditioning. We constructed a PV for all spatial bins of the three context A exposures (PV-A) and a second PV for all spatial bins of the three context B exposures (PV-B) using the rate maps of all recorded cells that excluded periods of immobility. Then, we correlated each spatial PV with the instantaneous neuronal activity (**Figure 3A**). As expected, context fear discrimination was associated with retrieval of spatial representations for contexts A and B across context presentations on day 3 in dCA1 and vCA1 (**Figure 3B-C**). Indeed, in dCA1 and vCA1 context A presentations produced a greater mean correlation with PV-A compared to PV-B, and conversely when context B was presented (**Figure 3B-C**). We analyzed the mean correlation values of the spatial PV corresponding to the presented context (i.e. correlation with PV-A when context A is presented; correlation with PV-B when context B is presented) and found that context representations were overall stronger in dCA1 compared to vCA1 (**Figure 3D; Figure S4E, S4F**). Furthermore, single cell remapping analyses indicated that dCA1 was more stable than vCA1 during day 3 context fear discrimination (**Figure S3**). To control for within-session fear extinction, we quantified the mean correlation of values of PV-A and PV-B only during the first two context exposures, i.e., from 0 to 4 min on day 3. As in the entire discrimination test **(Figure 3D)**, we found a stronger representation in dCA1 **(Figure 3E)**. To assess the magnitude of spatial representation discrimination, we calculated the within-subject difference of PV-A and PV-B correlations during the context presentations. We observed a stronger difference in dCA1 compared to vCA1 during context B presentation **(Figure 3E)**. Importantly, in vCA1 there was no significant difference in the PV-A correlation when contexts A and B were presented **(Figure 3C, right),** whereas the PV-A correlation significantly decreased during context B exposure for dCA1 **(Figure 3B, right).** Together, these results suggest that the vCA1 representation of the conditioned context remains strong even when the neutral context is presented. Thus, both dCA1 and vCA1 neuronal activities dynamically alternated according to the context displayed, but in dCA1 the representations of context A and context B were more distinct.

**Figure 3:**
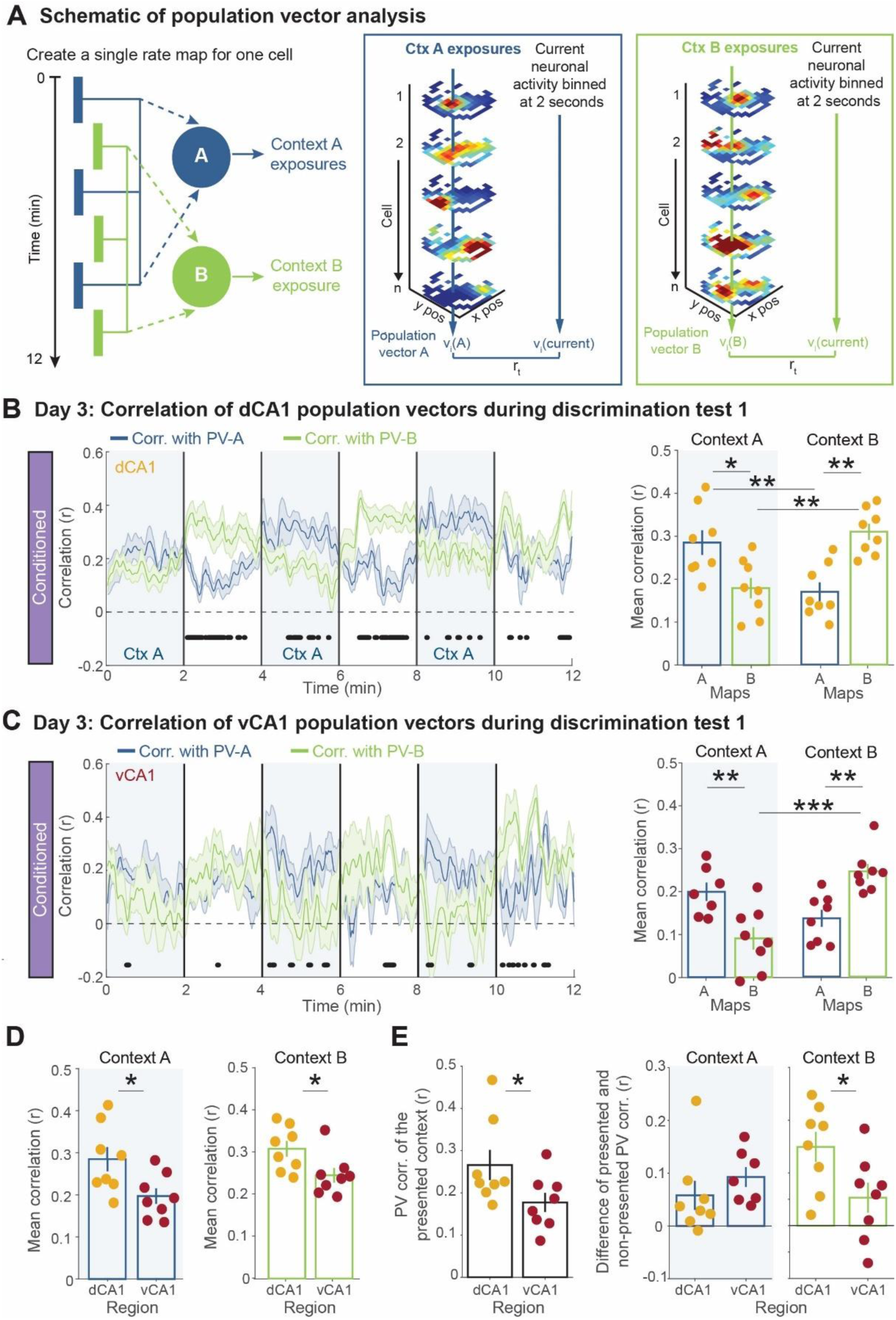
Context A and B spatial representations are more distinct in hippocampal dCA1 compared to vCA1 during the discrimination test. **(A)** Schematic of the population vector (PV) analysis method. Left, construction of a PV for all spatial bins of the three context A exposures (PV-A) and a second PV for all spatial bins of the three context B exposures (PV-B). PV-A and PV-B are correlated (Pearson’s r) with the current neuronal activity to generate PV correlations for each context, across time. **(B)** Left, dynamics of population vector correlations with PV-A or PV-B in dCA1 conditioned group (n = 8 mice) during day 3 discrimination test. Blue shaded regions indicate context A exposures, white regions are context B periods. Black dots indicate timepoints of significant pairwise difference between PV-A and PV-B correlation values (Wilcoxon signed-rank test, p < 0.05). Right, mean correlations with PV-A or with PV-B during contexts A and B exposures in dCA1 conditioned group shows that the average PV correlation that corresponds to the present context is greater than the PV correlation that corresponds to the non-presented context (context A Wilcoxon signed rank test: signed rank = 36, p < 0.01; context B Wilcoxon signed rank test: signed rank = 0, p < 0.01). **(C)** Left, dynamics of PV correlations with PV-A or PV-B during day 3 discrimination test as in **(B)** but recorded from vCA1 conditioned group (n = 8 mice). Right, mean correlations with PV-A or with PV-B during contexts A and B exposures in vCA1 as in (**B**). Significant comparisons in vCA1 conditioned group were similar to dCA1 in (**B**) (Context A: Wilcoxon signed rank test: signed rank = 36, p < 0.01; Context B: Wilcoxon signed rank test: signed rank = 0, p < 0.01). **(D)** Mean PV correlations of contexts A and B recorded in dCA1 (n = 8 mice) are significantly greater than mean PV correlations recorded in vCA1 (n = 8 mice) (Context A: Wilcoxon signed rank test: signed rank = 89, p < 0.05; Context B: Wilcoxon signed rank test: signed rank = 90, p < 0.05). **(E)** Analysis during the first context A and B presentations. Left, mean PV correlation of the presented context (PV-A during context A and PV-B during context B) in each mouse was significantly higher in dCA1 compared to vCA1 (Wilcoxon rank test: rank = 89, p < 0.05). Right, the difference between the mean PV correlation of the presented context and the mean PV correlation of the non-presented context was calculated to assess the magnitude of spatial representation discrimination. Similar results were found between dCA1 (n = 8 mice) and vCA1 (n = 7,8 mice; 1 vCA1 mouse excluded due to inadequate spatial sampling in context A) when presenting context A (Wilcoxon rank test: rank = 49, p > 0.05) but not during context B exposure (Wilcoxon rank test: rank = 87, p < 0.05). Data are expressed as the mean ± standard error of the mean.

### Context fear conditioning differently affects representations along the hippocampal longitudinal axis

To directly assess the impact of fear conditioning on spatial map similarity in the hippocampus, we compared the conditioned group to the non-conditioned control group. First, we confirmed that in the absence of conditioning, both dCA1 and vCA1 generate distinct PVs for contexts A and B during the day 3 discrimination test (**Figure 4A-D**). As expected in dCA1 and vCA1, the average PV correlation of the presented context was significantly greater than the PV correlation of the non-presented context (**Figure 4B, 4D**).

**Figure 4:**
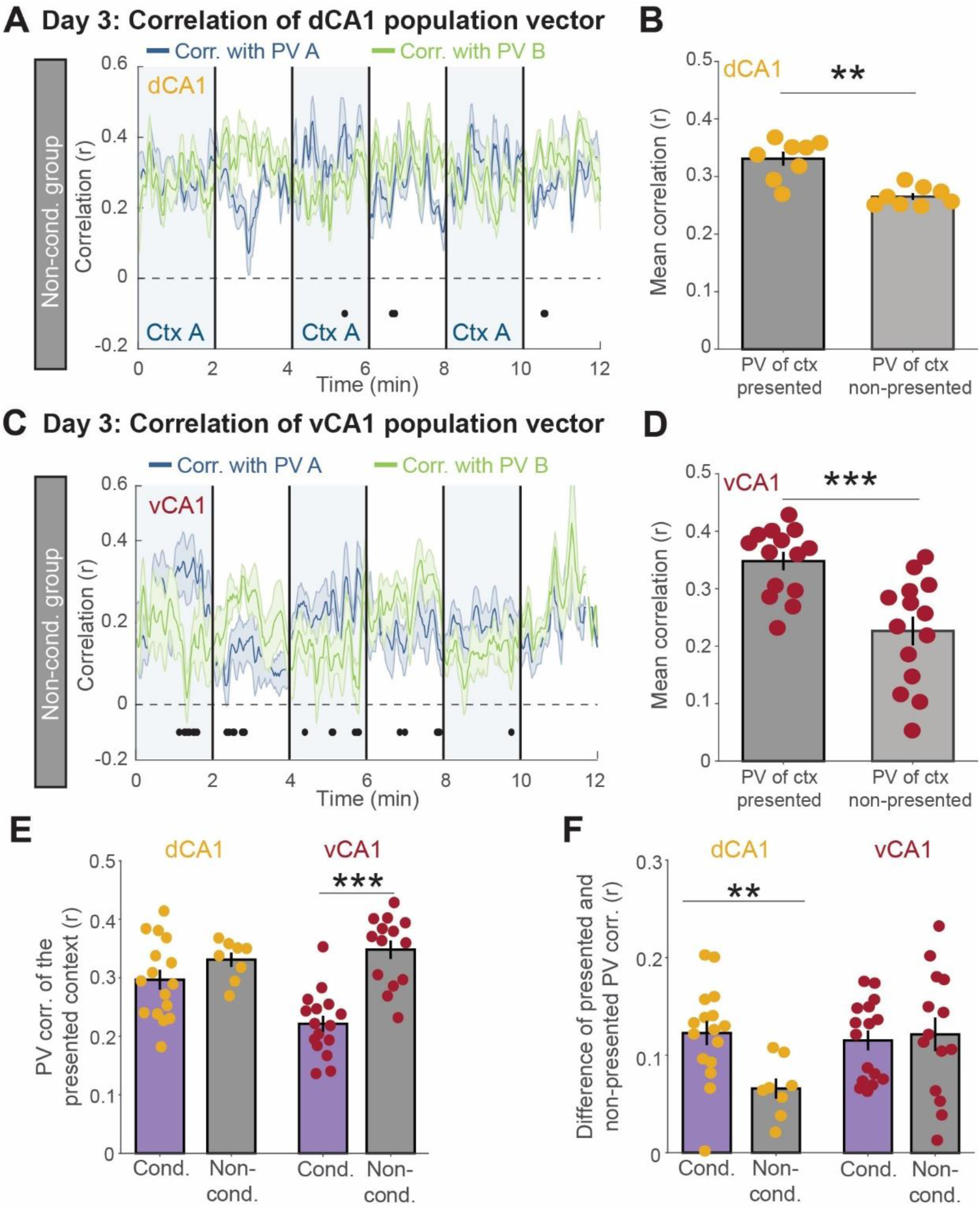
Context fear conditioning differentially affects spatial representations along the hippocampal longitudinal axis. **(A)** Dynamics of dCA1 population vector correlations with PV-A or PV-B in the non-conditioned group (n = 4 mice) during day 3 discrimination test. Blue shaded regions indicate context A exposures, white regions are context B. Black dots indicate timepoints of significant (Wilcoxon signed-rank test, p < 0.05) pairwise comparisons between PVs A and B correlation values. **(B)** Correlations with the PV of the context presented (either with PV-A during presentations of context A or with PV-B during presentations of context B) are significantly higher than the correlations with PV of the non-presented context (Wilcoxon signed rank test: signed rank = 36, p < 0.01). **(C)** Dynamics of population vector correlations with PV-A or PV-B during day 3 discrimination test as in (**B**) but recorded from vCA1 non-conditioned group (n = 8 mice). Black dots indicate timepoints of significant (Wilcoxon signed-rank test, p < 0.05) pairwise comparisons between PVs A and B correlation values. **(D)** Correlations with the PV of the context presented are significantly higher than the correlations with the PV of the non-presented context (Wilcoxon signed rank test: signed rank = 105, p < 0.001). **(E)** The mean PV correlation of the presented context is significantly lower in vCA1 conditioned mice compared to non-conditioned mice (right: Wilcoxon rank test: rank = 148, p < 0.0001), but not in dCA1 mice (left, Wilcoxon rank test: rank = 180, p > 0.05). **(F)** Difference between the mean PV correlation of the presented context and the mean PV correlation of the non-presented context was calculated to assess the magnitude of spatial representation discrimination. PV differences were significantly higher in conditioned mice compared to non-conditioned mice in dCA1 (left, Wilcoxon rank test: rank = 246, p < 0.01), but not vCA1 (right, Wilcoxon rank test: rank = 242, p > 0.05). Data are expressed as the mean ± standard error of the mean.

Moreover, comparisons between conditioned and non-conditioned mice showed that fear conditioning exaggerated the difference between PV-A and PV-B correlations in dCA1 during a context presentation (**Figure 4F**), although conditioning itself did not strengthen context A and B representations in dCA1 (**Figure 4E**). In contrast, in vCA1 conditioning did not exaggerate the difference between PV-A and PV-B correlations, as in dCA1 (**Figure 4F**). Yet, in vCA1 fear conditioning itself significantly weakened context A and B spatial representations (**Figure 4E**). Importantly, control analyses that matched neuronal sample sizes did not affect these differences between conditioned and non-conditioned groups (**Figure S4G, S4H**). These results demonstrate that dCA1 spatial representations were not altered by fear conditioning itself, instead fear conditioning increased the expression of the spatial map for the presented context. In contrast, fear conditioning in vCA1 altered the spatial representations by increasing their similarity, but expression of the spatial map for the presented context was unaffected by fear conditioning. These results are supported by the finding that dCA1 neurons encoded more spatial information (**Figure S2C-D**). This dCA1-vCA1 dissociation reveals that fear conditioning engages different context encoding schemes throughout the longitudinal axis by varying context representational similarity. This result was unexpected because the predicted consequence of context representation similarity in vCA1, is reduced specificity of spatial map expression during context memory retrieval, but this was not the case.

### Context transitions reveal that conditioned and non-conditioned context representations in dCA1 and vCA1 predict freezing behavior

Next we investigated the temporal link between the retrieval of spatial representations and fear behavior. We calculated the within-subject difference between PV-A and PV-B over time and correlated these spatial PV differences with freezing behavior (**Figure 5A-B**). In both dCA1 and vCA1 we observed that strong correlations to PV-A, and weak correlations to PV-B resulted in more freezing behavior on day 3 and 4 discrimination tests (**Figure 5A-B**). During context transitions, we observed that PV-A had a stronger representation relative to PV-B after the transition to context A (**Figure 5C).** Conversely, transitions to context B were associated with rapid spatial representation switching from PV-A to PV-B (**Figure 5D**). These dynamics were strongly correlated with an increase in fear behavior when transitioning to context A (**Figure 5C, right**) or with a lower freezing behavior when transitioning to context B (**Figure 5D, right**). Taken together, these data indicate that the population dynamics during context teleportation in dCA1 and vCA1 predict global changes in freezing behavior.

**Figure 5:**
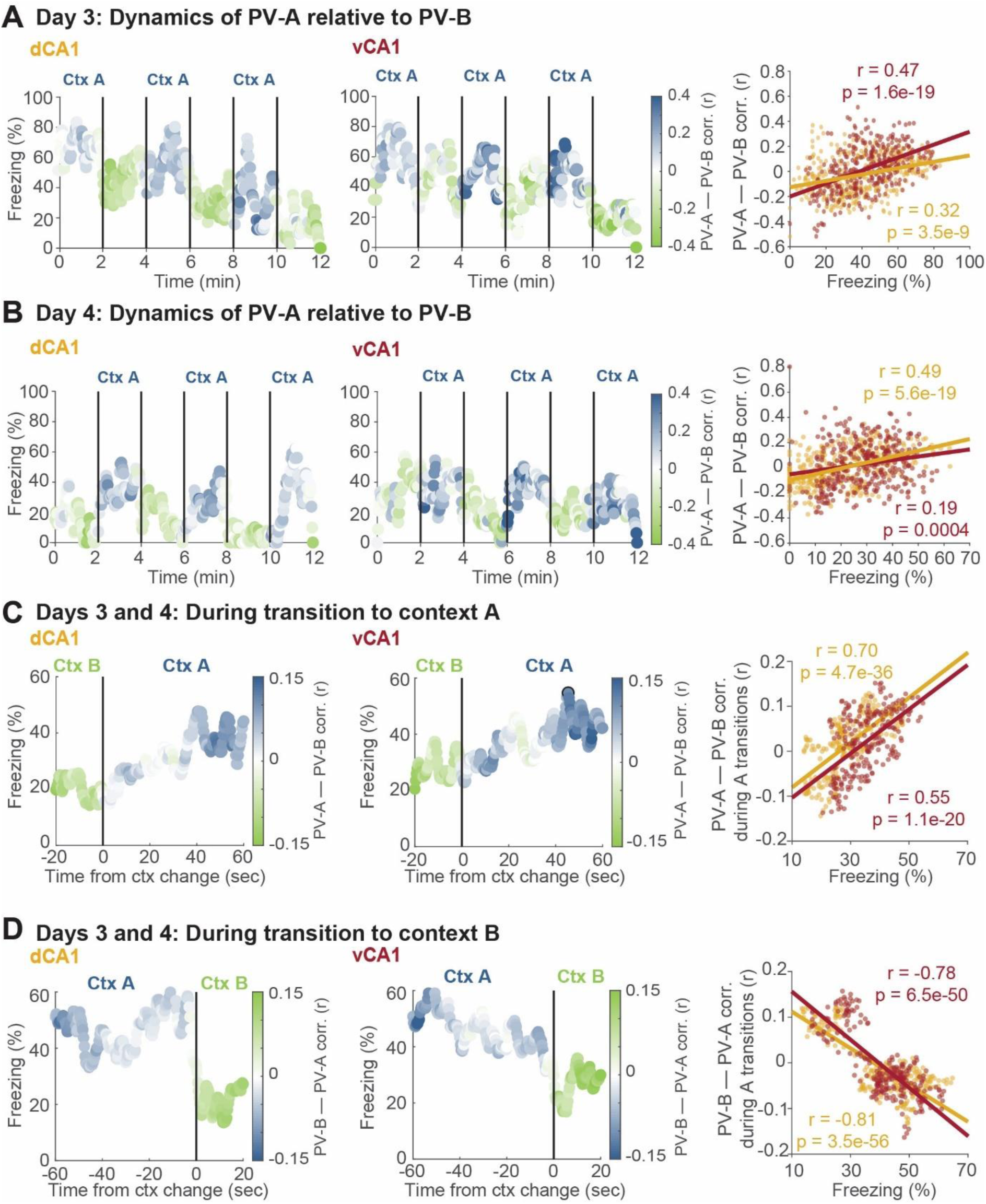
Context transitions reveal that differences in representations of conditioned and non-conditioned contexts predict freezing behavior. **(A)** Dynamics of PV-A relative to PV-B during discrimination day 3 in conditioned mice recorded in dCA1 (left, n = 8 mice) or in vCA1 (middle, n = 8 mice). Right, PV differences were calculated to assess magnitude of representational discrimination. Differences in PVs were significantly correlated with freezing behavior in both dCA1 (correlation coefficient r = 0.32, p < 0.0001, n = 360 time points) and vCA1 (correlation coefficient r = 0.47, p < 0.0001, n = 360 time points). **(B)** Left and middle, dynamics of PV-A relative to PV-B as in (**A**) but during discrimination day 4. Right, PV differences throughout test day 4 and freezing behavior is significantly correlated in both dCA1 (correlation coefficient r = 0.49, p < 0.0001, n = 360 time points) and vCA1 (correlation coefficient r = 0.19, p < 0.001, n = 360 time points). **(C)** Dynamics of PV-A relative to PV-B before and after context A presentation during days 3 and 4 discrimination tests for mice recorded in dCA1 (left) or in vCA1 (middle). Weaker PV-B and stronger PV-A correlations are associated with higher percent of freezing in both dCA1 (correlation coefficient r = 0.70, p < 0.0001, n = 241 time points) and vCA1 (correlation coefficient r = 0.55, p < 0.0001, n = 241 time points). **(D)** Dynamics of PV-B relative to PV-A before and after context B presentation during days 3 and 4 discrimination tests for mice recorded in dCA1 (left) or in vCA1 (middle). Weaker PV-A and stronger PV-B correlations are associated with lower probability of freezing in both dCA1 (correlation coefficient r = −0.81, p < 0.0001, n = 241 time points) and vCA1 (correlation coefficient r = −0.78, p < 0.0001, n = 241 time points).

### Fear extinction reverses fear conditioning-induced representational similarity in vCA1

Finally, we investigated freezing behavior in context A across days 1 to 4. In comparison to pre-shock freezing on day 1, freezing levels significantly increased during conditioning on days 1 and 2. In comparison to the first context A presentation on day 3, freezing behaviour was significantly reduced during repeated context A exposures on days 3 and 4 (**Figure 6A**).

**Figure 6:**
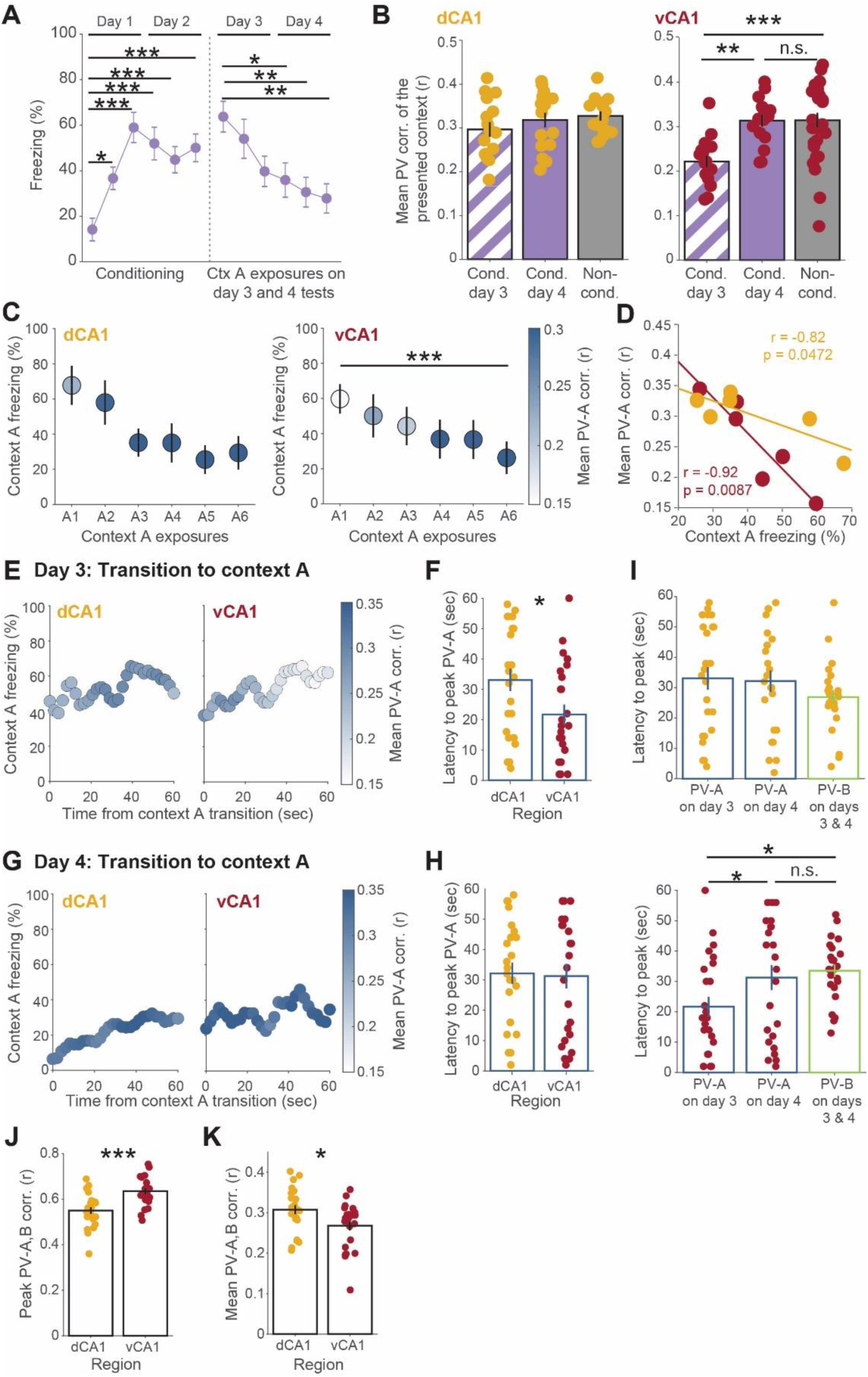
Fear extinction reverses fear conditioning-induced representational similarity in vCA1. **(A)** Left, fear conditioning on days 1 and 2 significantly increased freezing in dCA1 and vCA1 groups (n = 16 mice, one-way RM ANOVA: main effect of time F_(5,15)_ = 11.73, p < 0.0001 followed by a Bonferroni correction test). Right, fear extinction during context A exposures on discrimination test days 3 and 4 (one-way RM ANOVA: main effect of time F_(5,15)_ = 4.777, p = 0.0008 followed by a Bonferroni correction test). Statistical comparisons in the left panel are in reference to the first 4 min (pre-shock period) and in the right panel comparisons are to the first context A exposure. **(B)** Left, mean PV-A and PV-B correlations when contexts A and B are presented, respectively. No significant differences in conditioned (Cond.) mice on day 3 or day 4 (n = 8) and combined non-conditioned (Non-cond.) mice on days 3 and 4 (n = 4) in dCA1 (Kruskal-Wallis test: rank = 1.487, p = 0.4755). Right, mean correlation values were significantly lower in the conditioned group on day 3 compared to day 4 and combined non-conditioned mice on days 3 and 4 in vCA1 (Kruskal-Wallis test: rank = 16.67, p = 0.0002 followed by a Dunn comparison test). The statistics presented correspond to the Dunn comparison test. **(C)** Mean PV-A correlation and freezing behavior during context A presentations on days 3 (A1-A2-A3) and 4 (A4-A5-A6) for dCA1 (left, n = 8 mice) and vCA1 (right, n = 8 mice). The mean PV-A correlation decreased during subsequent context A exposures in vCA1, but not dCA1 (dCA1: Friedman’s test: F = 6.4, p > 0.05; vCA1: Friedman’s test: F = 21.6, p < 0.001). **(D)** Relationship between the mean PV-A correlation values and freezing behavior in context A (dCA1 correlation coefficient r = −0.82, p < 0.05, n = 6 context A exposures; vCA1 correlation coefficient r = −0.92, p < 0.01, n = 6 context A exposures). **(E)** Dynamics of PV-A after transition to context A in 2 sec intervals during day 3 discrimination test in conditioned mice recorded in dCA1 (left) or in vCA1 (right). **(F)** Latency to the peak PV-A during transitions to context A on day 3 (n = 8 mice for 3 context A transitions, Wilcoxon rank test: rank = 693, p < 0.05). **(G)** Dynamics of PV-A correlation from transition to context A as in (**E)** but during day 4 discrimination test. **(H)** Latency to the peak PV-A during transitions to context A on day 4 (n = 8 mice for 3 context A exposures, Wilcoxon rank test: rank = 590, p > 0.05). **(I)** Top, no significant difference in dCA1 latency to peak PV-A or PV-B correlations during context transitions on days 3 and 4 (Kruskal-Wallis test: rank = 2.291, p = 0.32). Bottom, in vCA1 the latency to the peak PV-A correlation during transitions to context A was significantly lower during discrimination day 3 compared to discrimination on day 4 and compared to PV-B during transitions to context B on days 3 and 4 (Kruskal-Wallis test: rank = 6.758, p = 0.03 followed by a Dunn comparison test). The statistics presented correspond to the Dunn comparison test. **(J)** Peak values of PV-A or PV-B correlations during transition to their corresponding context on days 3 and 4 were significantly higher in vCA1 compared to dCA1 (Wilcoxon rank test: rank = 400, p < 0.001). **(K)** Mean of PV-A or PV-B correlations during the entire time of their corresponding context on days 3 and 4 were significantly higher in dCA1 compared to vCA1 (Wilcoxon rank test: rank = 709, p = 0.01). All data are expressed as the mean ± standard error of the mean.

We then examined the time course of hippocampal representations on day 4 discrimination test (**Figure S5**) and as expected, we observed that the average correlation of PVs during the presented context was higher than the non-presented context in both dCA1 and vCA1 (**Figure S5B, S5G**). While dCA1 sustained a robust PV correlation of the presented contexts on days 3 and 4, in vCA1 the PV correlations of the presented contexts were augmented on day 4 to values observed in non-conditioned mice (**Figure 6B**). This surprising effect indicated that fear extinction reversed the fear conditioning-induced representational similarity of PV-A and PV-B in vCA1 (**Figure 4E**). To explore this further, we calculated the mean PV-A correlation across context A exposures on days 3 and 4 and plotted it against freezing behavior. Although both vCA1 and dCA1 showed similar freezing behavior reduction, only vCA1 had significant increases in PV-A correlation with successive context A exposures (**Figure 6C**). We observed a strong negative association between PV-A correlations and freezing behavior in vCA1, this association was much weaker in dCA1 (**Figure 6D**). Moreover, this association between vCA1 spatial representation strength and freezing reduction was robust, as it was replicated with simply PV-B, or when PV-A and PV-B were considered in parallel (**Figure S6A-E**).

We then examined context A spatial representations during context A transitions on day 3 and observed that both dCA1 and vCA1 displayed increasing PV-A correlations following transition (**Figure 6E**). However, dCA1 and vCA1 differed markedly in (1) the relationship between the PV-A correlation and freezing behavior and (2) the temporal dynamics of the PV-A correlation value. We observed that the latency to peak PV-A correlation value was significantly shorter in vCA1 compared to dCA1 during context A transitions (**Figure 6F**), suggesting a time-specific role for the vCA1 population representation at the moment of context memory retrieval. Conversely, during transitions to neutral context B, no significant differences in PV-B peak latencies were observed between dCA1 and vCA1, though vCA1 latencies tended to be longer compared to dCA1 **(Figure S6F-G and S6I-J)**. Thus, vCA1 spatial representations of the fear conditioned context are most strongly expressed immediately following the retrieval of the fear conditioned context memory on day 3.

However, on day 4 when context fear extinction is evident (**Figure 6A**), we observed that in vCA1 the relationship between context A representation retrieval and fear behavior became similar to dCA1 in magnitude, and temporal dynamics. Additionally, the relationship between context A spatial representation and freezing behavior in vCA1 was no longer present (**Figure 6G**). Importantly, during fear extinction on day 4 the latencies to maximum PV-A correlation became nearly identical in vCA1 and dCA1 **(Figure 6H)**. Indeed, we observed an increased latency for PV-A to reach peak correlation on day 4, producing latencies similar to those observed with PV-B in vCA1, whereas no difference in peak PV latencies were observed in dCA1 **(Figure 6I)**. Although vCA1 PV-A latencies normalized to the level of PV-B, the maximum correlation values within 60 seconds of context A or B transition were higher in vCA1 on days 3 and 4 (**Figure 6J, Figure S6H, S6K**). However, as previously observed (**Figure 3E**) the mean PV correlations calculated during the entire context exposure was lower in vCA1 compared to dCA1 (**Figure 6K**). These results indicate that vCA1 retrieval dynamics of a threatening context are maximally expressed prior to the stable spatial representation observed in dCA1. Together these findings reveal a temporal dissociation between vCA1 and dCA1 context representations during emotional memory retrieval, wherein vCA1 initially generates a strong spatial representation, followed by a slower, but more stable dCA1 spatial representation.

## Discussion

The hippocampus is well-known for its spatial and context-dependent representations ^5,9,30^, yet the link between these representations and memory acquisition, retrieval, and extinction have been difficult to resolve due to technical limitations. Here we combined a novel context teleportation protocol with 1-photon calcium imaging in dCA1 and vCA1 to record and track large populations of neurons across multiple phases of a context fear memory. We found that fear memory encoding most rapidly and robustly changed vCA1 representations, however, to our surprise the representations of the neutral and conditioned contexts became more similar in vCA1. The fear conditioning-induced representational similarity in vCA1 was accompanied by rapid switching between these context representations and more strongly predicted freezing behavior, compared to dCA1. When fear extinction reduced the threat-level of the context, vCA1 representations returned to states observed prior to fear conditioning. These results indicate that vCA1 spatial representation retrieval dynamics are tightly matched to the presence of threat in a context. Together our findings reveal that vCA1 has a privileged role in threat memory acquisition, expression, and extinction.

### Development of a context memory retrieval task

We developed an apparatus that permitted us to manipulate the context while the mouse remained undisturbed in the same physical space. Two previous studies developed tasks that fulfilled these requirements in freely behaving mice and revealed the moment-to-moment alternations in hippocampal and medial prefrontal activity during context transitions. Jezek et al 2011 ^14^ used an apparatus with differential lighting to teleport rats between two contexts while foraging for food and they found dorsal CA3 momentarily *flickered* between context representations. In a second study by Rozeske et al 2018 ^52^, researchers manipulated visual, auditory, and olfactory stimuli during a context fear discrimination task and observed distinct cortical representations of the conditioned and neutral contexts. Here, we used a large context and manipulated visual and auditory stimuli to investigate hippocampal representations during teleportation between conditioned and neutral contexts. Our task permitted investigation of the dynamic relationship between fear memory and alterations in dCA1 and vCA1 context representations. The present results replicate previous work ^14^ that found context teleportation alters hippocampal context representations and establishes a new link between hippocampal context representations and context-guided behavior. Our findings demonstrate that our task effectively teleports mice and produces retrieval of a threatening memory, allowing us to directly examine the relationship between cognitive mapping and memory retrieval.

### Functional differences in spatial and emotional processing along the hippocampal longitudinal axis

The dorsal and ventral CA1 are functionally distinct structures ^31,32,56,57^ with variations in anatomical connectivity ^58^, gene expression ^59^, cell excitability ^60^, and morphological properties ^60,61^. Given this diversity, it is not surprising that the dorsal hippocampus functionally supports spatial representations whereas the ventral hippocampus is implicated in emotional processing ^31,32,56,62,63^. Yet, some studies reported that the ventral hippocampus also has a key role in various forms of learning including goal-oriented searching, conditioned inhibition, spatial learning, and discriminative fear conditioning to similar contexts ^64,65^. Our apparatus was designed to combine spatial and emotional task features so we could dissociate functional contributions of dCA1 and vCA1. We observed both regions were engaged in all phases of testing, but vCA1 was most sensitive to threatening stimuli. Although dCA1 and vCA1 recordings were in separate mice, statistical comparisons were possible because the number of mice and fear behavior were nearly identical between groups. Moreover, despite different neuronal yields between groups, we sub-sampled dCA1 neurons in multiple control analyses to match vCA1 cell numbers and found our spatial representation differences between dCA1 and vCA1 were robust. Current technology does not permit dual miniscope recordings in dCA1 and vCA1, but future studies should examine how spatial and emotional representations interact during fear memory acquisition, retrieval, and extinction. Together, our results are consistent with previous findings that dissociate spatial and emotional functions along the hippocampal longitudinal axis ^31,32,56,62,63^.

### Context representation updating during threat memory acquisition

We observed that vCA1 context representation was strongly influenced by footshock delivery compared to dCA1. Moreover, in vCA1, but not dCA1, there was a strong positive correlation between the magnitude of spatial representation change during fear conditioning and fear behavior expression. Other reports align with our finding that vCA1 is sensitive to threatening stimuli. These include a necessary role of vCA1 in context fear conditioning ^49^, defensive behaviors in response to innate predator odor ^66^, anxiety ^37,67,68^, preferential projections from the ventral hippocampus to “fear neurons” in the basolateral amygdala ^69^, and active avoidance responses ^45^.

The current study used miniscope imaging to record dozens of spatially tuned cells to construct spatially-rich population vectors for each mouse. Although previous electrophysiology recordings in dCA1 observed a diversity of place cell responses following threatening stimuli ^9,30,70,71^, cell yields did not permit construction of context population vectors in all subjects. Miniscope imaging allowed us to assess for the first time the population dynamics of a context representation, possibly the cognitive map, in each mouse during threat memory encoding. The vCA1 representation changes during fear conditioning are likely supported by cells that respond to shock and are essential for contextual fear memory ^43^. Moreover, the rapid switching of context representations in vCA1 and the strong relationship between context representation and freezing behavior, suggests that vCA1 is highly sensitive to threatening stimuli, or perhaps more generally context valence. Previous work demonstrates that altering reward contingencies in a context rapidly remaps place cells in the intermediate, but not dorsal, hippocampus ^72^. Overall, our results demonstrate that vCA1 predominately represents threatening stimuli and may exclusively orchestrate threat-related behavior.

### Context representation specificity and dynamics during fear memory retrieval

Here we linked the hippocampal spatial representation of a context memory with the expression of a threat memory. We observed that in both dCA1 and vCA1 the strength of the PV-A did not reliably predict freezing behavior. Rather, the difference between PV-A and PV-B at a given moment during our context discrimination task most reliably predicted freezing behavior. These results suggest that when confronted with somewhat similar contexts, the hippocampus may continuously multiplex competing context representations. We found that the moments wherein one representation dominated over the other are the periods that most reliably predicted context-guided behavior, i.e. freezing or not. These moments of representational switching were most pronounced in vCA1, where the context representational similarity was high compared to dCA1. Our results indicate that the representational similarity of differing contexts within vCA1 may provide a rapid, low resolution, gist representation of the context, whereas the dCA1 supports a highly detailed cognitive map ^38,73,74^. Our results also support the possibility that vCA1 has a unique population coding scheme to provide rapid and efficient switching between context representations. Supporting this possibility, we observed that vCA1 context population codes for neutral and threatening contexts became more similar after fear conditioning, and during context teleportation the transition to the new network state was faster. Previous modeling work supports the idea that transitions between ‘shallow’ basins in an attractor network, which can arise from an overlapping population code, would support these faster state transitions ^75^.

Due to the relatively slow kinetics of GCaMP6f, our study was not able to examine the mechanism of representational cycling ^76,77^, but one possibility is that within the trisynaptic circuit there is a continuous cycle of pattern completion and separation processes, rather than a singular moment of pattern completion. Indeed in a previous context teleportation study ^14^, context representations momentarily flickered at the theta rhythm prior to pattern completing to one of the context representations. Our calcium imaging recording method does not provide the temporal resolution to assess theta-rate flickering, but our findings are in agreement with transient periods context representation.

This study also suggests that the behavioral expression of a memory does not appear to be a direct consequence of context representation retrieval. We observed that context fear conditioning altered the hippocampal spatial representation in real time, despite the stability of the physical context. Moreover, during context teleportation we observed replicable, within-subject, context representation retrievals. These results demonstrate that the spatial representation is a reliable substrate for the memory as it is modified by the conditioning experience and is strongly expressed during presentation of the conditioning context. Yet, we observed a dissociation of the context representation memory and the behavior-based memory. Indeed, the strength PV-A after teleportation to context A did not reliably predict fear behavior in either vCA1 or dCA1. If context representations are sufficient for behavioral expression of the memory, then moments of robust context representation would precisely coincide temporally with freezing behavior. Yet, freezing epochs were not temporally tied to the strength of the hippocampal spatial representation. This finding suggests a downstream structure, presumably the basolateral amygdala ^69^ or medial prefrontal cortex receives weighted context representations and initiates freezing behavior accordingly.

### Effect of context fear extinction in spatial representation along the hippocampal longitudinal axis

We observed that vCA1 representations were highly correlated with the threat-level of the context and fear expression. Following context fear conditioning, vCA1 representations were drastically altered compared to non-conditioned controls, but following fear extinction, vCA1 spatial representations returned to levels comparable to vCA1 control mice. Spatial representations in conditioned and non-conditioned dCA1 groups were not significantly altered throughout the task. Theoretical and empirical work suggest that fear extinction is either “unlearning” or a form of new learning in which an association between the conditioned stimulus and absence of the unconditioned stimulus is formed ^23^. Although there are no vCA1 recordings during pure context fear extinction (i.e. tests that did not include foreground conditioning of an auditory tone with footshock), some studies have examined dCA1 activity during context fear extinction. Immediate early gene staining in a within-subject design revealed that largely separate populations of dCA1 neurons were labeled following context fear conditioning and extinction^78^. Corroborating the existence of a distinct hippocampal cell population for context fear acquisition and extinction is an activity-dependent tagging study in the dentate gyrus ^79^. The authors optogenetically inhibited the dentate gyrus to demonstrate that fear acquisition and extinction neurons are necessary to either express, or prevent, freezing behavior.

Previous studies examining fear extinction that used *in vivo* recording techniques only examined the dorsal hippocampus and indicate that a new extinction neural representation is not consistently observed. During a non-spatial trace fear conditioning task, calcium imaging in dCA1 neurons revealed distinct populations for acquisition and extinction ^80^. In contrast, 2-photon calcium imaging of synapses strengthened between CA3 and CA1 during context fear memory acquisition become uncoupled during fear extinction^81^. Moreover, dCA1 place cell recordings during extinction to a predator odor revealed only approximately 20% of previously stable place cells remapped^82^. The authors note that place cells did not display long-term stability after extinction and speculated that place cells may in fact revert to their previous place fields. Our population vector analysis in vCA1 is the first clear demonstration that following context fear extinction, the context representation returns to the original state. This context representation reversal may be more exaggerated in vCA1 compared to other hippocampus regions and for these reasons previous literature is not uniform.

## Conclusion

Contextual fear conditioning uniquely depends on both spatial and emotional processing in the hippocampus. The present study assessed the contributions of functionally distinct dCA1 and vCA1 regions during encoding, retrieval, and extinction of context fear memory. Our recordings demonstrate that spatial representations in vCA1 become more similar following fear conditioning and generate faster threat memory retrieval compared to dCA1. Conversely, when context threat was reduced by fear extinction, vCA1 context representations returned to their pre-conditioning state. Together our results demonstrate that vCA1 network activity is intimately influenced by threatening memories and suggest that vCA1 contains a unique network coding scheme for rapid and efficient switching between emotional states.

## Acknowledgments

We thank D. Aharoni and the miniscope.org community for sharing their knowledge and expertise as well as G. Lopez for technical assistance with *Bonsai* software. We are grateful to J.Q. Lee, C-A. Mosser, M. Oulé, M.H. Yaghoubi, and H. Nagaraj for providing helpful comments on prior versions of the manuscript, as well as to all members of the Brandon Laboratory for useful discussions. This research was funded by the Canadian Institutes of Health Research (CIHR project grants #367017 and #377074), the Natural Sciences and Engineering and Research Council of Canada (NSERC Discovery grant #74105), and the Canada Research Chairs Program awarded to M.P.B. This work was also supported by CIHR fellowship (#158096) to R.R.R. L.R. is supported by a Doctoral Training fellowship, the Shuk-Tak Liang Fellowship (Faculty of Medicine Internal Studentship).

## Author contributions

R.R.R., L.R. and M.P.B. conceived the project.

R.R.R., L.R. and A.S. developed the fear discrimination task.

R.R.R. and L.R. conducted all experiments.

R.R.R., L.R., A.T.K. wrote codes for analysis.

R.R.R. and L.R. performed data analysis.

R.R.R., L.R. and M.P.B. wrote the manuscript.

## Declaration of interests

Authors declare no competing interests

## Methods

### RESOURCE AVAILABILITY

#### Lead contact

Further information and requests for resources should be directed to and will be fulfilled by the lead contact, Mark P. Brandon.

#### Materials availability

This study did not generate new unique reagents.

#### Data and code availability

All data reported in this paper will be shared by the lead contact upon request. All original code will be deposited at GitHub and is publicly available as of the date of publication DOIs are listed in the key resources table. Any additional information required to reanalyze the data reported in this paper is available from the lead contact upon request.

### EXPERIMENTAL MODEL AND SUBJECT DETAILS

#### Experimental animals

Male C57BL/6J wildtype mice (8-10 weeks old) were obtained from Charles River Laboratory. Mice were individually housed in a temperature- and humidity-controlled animal facility (21 ± 2 °C, 45 ± 5% humidity) on a 12 hour dark-light cycle with food and water *ad libitum*. Experiments were conducted during the light cycle.

All animal experiments were approved by the Facility Animal Care Committee (FACC) of McGill University and carried out in accordance with the Douglas Hospital Research Centre Animal Use and Care Committee (protocol #2015-7725) and Canadian Institutes of Health Research guidelines.

### METHOD DETAILS

#### Viral construct

AAV5.CaMKII.GCaMP6f.WPRE.SV40 was the adeno-associated virus (AAV) used for calcium imaging and was obtained from Addgene at 2.3e13 GC-ml. Prior to surgical microinjection it was diluted 1:1 in sterile PBS.

#### Surgeries

Mice were anesthetized with isoflurane (4.5% for induction and 2% for maintenance during surgery) in oxygen. Mice were placed in a stereotaxic frame (David Kopf Instruments) and body temperature was stabilized with a heating pad. Analgesic Carprofen at 20 mg/ml (10 ml/kg) and 0.9% saline solution (10 ml/kg) were subcutaneously injected, and eyes were hydrated with gel (Optixcare).

For miniscope imaging experiments, mice were unilaterally microinjected with 400-600 nl of AAV to express GCaMP6f (Nanoject II; Drummond Scientific) at the following coordinates (mm) relative to dura: dCA1 −2.00 AP, 1.75 ML, −1.4 DV; vCA1 −3.16 AP, 3.25 ML, −3.85, −3.50, −3.25 DV. Mice were allowed at least three weeks to recover from surgery. During a second surgery under isoflurane anesthesia dCA1 mice were stereotaxically implanted with a 1.0 mm diameter gradient refractive index (GRIN) lens (Go!Foton) over the injection site after intervening cortical tissue was aspirated. In vCA1, a 0.5 mm diameter GRIN lens (Inscopix) was slowly lowered to the target depth (in mm from skull, vCA1: −3.16 AP, 3.50 ML, −3.50 DV). Two self-tapping screws were affixed to the skull and GRIN lenses were secured with dental cement (C&B Metabond). A silicone adhesive (Kwik-Sil, World Precision Instruments) was applied to protect the surface of the lens until the baseplating procedure. At least two weeks after lens implantation, an aluminum baseplate was affixed with dental cement to the existing dental cement from the GRIN lens implantation under isoflurane anesthesia. Mice were allowed at least one week to recover. After all surgeries, mice were placed on a heating pad and continuously monitored until they were mobile, then returned to their homecage.

#### Context fear retrieval task

The apparatus was composed of a large cylindrical LED screen (77 cm high, 90 cm diameter; Shenzhen Apexls Optoelectronic Co.) that controlled the visual context. The LED screen was placed on a large (92 cm x 92 cm) stainless-steel grid (Imetronic) to deliver scrambled footshocks (Coulbourn Precision Animal Shocker). Auditory stimuli were delivered via speakers mounted below the grid floor. Finally, an array of white LED lights (Luminosum) were located below the shock grid floor. The apparatus was controlled by a PC equipped with Bonsai software^83^ and a PulsePal (Sanworks LLC).

Mice were habituated to the apparatus in context C (black walls, flat white flooring, no sound, 1 lux illumination) for 10 min one day prior to experimentation. On days 1 and 2, mice were placed in context B (gray projected on the walls, white LED on flooring, 10 kHz tonic pure tone at 85 dB, 3 lux total illumination in the apparatus) for 12 min. Six hours later mice were placed in context A (Marshall Islands flag projected on the walls, unlit flooring, white noise at 75 dB, 73 lux total illumination in the apparatus) and six scrambled footshocks (0.6 mA, 1 sec) were delivered pseudorandomly during the 12 min context exposure. Between the two context exposures, mice were placed in their homecage and the apparatus was cleaned with 70% ethanol. On day 3, mice were tested in successive A and B contexts with an instantaneous transition, or *teleportation*, between the contexts at two minute intervals (configuration ABABAB). Day 4 was similar to day 3, except mice were tested in configuration BABABA (Figure 1 A-C). We included a non-conditioned control group that received the same testing protocol, except no footshock was delivered. Inclusion of these control groups is notable for two reasons: (a) experiments using 1-photon imaging during behavior often do not use a separate control groups because of the method is technically difficult and time consuming and (b) hippocampal representation changes could be assessed and compared within (context A vs B) and between (conditioning and non-conditioned) subjects, elevating the overall rigor of our experiments and validation of our teleportation task.

#### Histology and immunostaining

##### Perfusion

Mice were euthanized with isoflurane and transcardially perfused with 4% PFA in 0.01M PBS. Mouse brains were dissected out and postfixed in 4% PFA at 4°C overnight. The brains were transferred to a 0.01M PBS solution containing 30% sucrose and 0.1% sodium azide. Brains were sectioned into 40 μm coronal slices with a vibratome (VT1200, Leica Biosystems). Slices were stored in 0.01M PBS at 4°C until immunohistochemical processing.

##### Immunohistochemistry

Slides were blocked in a 0.01M PBS solution containing 0.25% Triton X-100 and 0.45% gelatine, three times during 15 min at room temperature. Staining was performed using 1:1000 primary antibody: rabbit anti-GFP (Life Technologies, Cat. No. A-11122) in blocking solution at 4°C overnight. After three washes, secondary antibody: donkey anti-rabbit (Molecular Probes, Cat. No. A-21206) were incubated 2 hours at room temperature. Slices were mounted on glass microscope slides (Adhesion Superfrost Plus Glass Slides, Brain Research Laboratories) and cover-slipped with a DAPI-containing mounting media (Fluoromount G with DAPI, eBioscience).

##### Slides scanner imaging

Fluorescent images from brain tissue were acquired with a slide scanner (Olympus VS120) and analyzed with ImageJ software (NIH).

#### Data acquisition

##### Locomotion

Locomotor behavior was recorded at a rate of 30 FPS using the Logitech c930e camera mounted to the ceiling of the apparatus. Video acquisition was initiated with Bonsai software^83^ and coordinated with calcium imaging with the miniscope data acquisition box (DAQ).

##### Miniscope calcium recording

As previously described^55^, dCA1 and vCA1 pyramidal cell activity was recorded with a miniscope connected to a DAQ (v3; miniscope.org) with a lightweight, flexible coaxial cable (Cooner Wire). Calcium activity was acquired at 30 Hz and imaged using an excitation LED set between 4–20%, depending on signal quality and the minimum possible light intensity to mitigate photobleaching. The gain was also adjusted on a mouse-by-mouse basis to avoid fluorescence signal saturation. Calcium imaging data was acquired with miniscope acquisition software (miniscope.org) and initiated by Bonsai software that controlled the behavioral task.

### QUANTIFICATION AND STATISTICAL ANALYSIS

#### Fear behavior

The hue-saturation-value (HSV) was modified in behavior videos to isolate the mouse from the surroundings and calculate the centroid position of the mouse for each video frame. Post-processing of the mouse’s trajectory included linear interpolation for the occasionally missed tracking frames and then was smoothed with a sliding window of 0.462 sec. Instantaneous speed was calculated and smoothed with a sliding window of 0.066 sec. A freezing epoch was defined as a speed less than 1 cm/sec, for at least 2 sec. These criteria most accurately described when the mouse had no visible movements and adopted typical freezing behavior.

#### Miniscope calcium recording

Calcium imaging data were preprocessed with a pipeline of open source MATLAB (MathWorks; version R2015a) functions^55^ to correct miniscope field of view movement using NoRMCorre^84^ and to segment putative cells and extract calcium transients using CNMF-E^85,86^. Spike likelihoods were extracted from calcium transient ΔF/F using a deconvolution algorithm. All analyses used a binarization method that treated the top 30% of spike likelihoods as inferred likelihood of spiking events (ILSE)^87^ and neuronal spike trains were constructed (Figure 2B). The signal-to-noise ratio (SNR) for each ΔF/F trace was calculated for each session as a measure of data quality as previously described ^55,88^.

To optimize recording of spatially-tuned cells in the hippocampus we expressed a calcium indicator under the control of CaMKII. In the hippocampus, CaMKII is expressed in pyramidal cells ^66,89^, which is the cell type with place cell functional properties ^90,91^. Therefore, all recorded cells were included in all spatial rate map analyses. Rate maps were constructed with bins of 7 cm and smoothed with a Gaussian Kernel of 10 cm. To quantify spatial information content of hippocampal cells ^55,92^, the rate maps from the first exposure to neutral context B were used.

For remapping analysis, we subsampled our data to match the spatial sampling distributions across all comparisons to correct for biases in sampled spatial locations (between pre- and post- conditioning or between contexts A and B exposures), as previously described^55^. We also only considered non-freezing periods. For the correlation between two same contexts, we compared the first half of the exposure to this context with the second half. Place field stability was determined by correlating place field maps (Pearson’s r) during first and second halves of recording; if a place field had less than 6 bins after subsampling it was not considered in the analysis.

Our general approach to understand spatial representations in the hippocampus was to construct a population vector (PV) by concatenating the spatial bin values of all rate maps in a mouse. Freezing periods were not included when constructing PVs. Specifically, on the conditioning day we constructed a PV that only considered spatial bins visited during the pre-shock period to create a pre-shock spatial representation of the context. For the discrimination tests, we constructed one PV that contained spatial bins visited during context A exposures (PV-A) and another one for spatial bins visited during context B exposures (PV-B). Then, we binned the PV at 2 sec and applied a smoothing window of 6 sec. During context transition periods when greater temporal precision was required the PV was binned at 0.33 sec. Next, to explore if experimental manipulations altered the spatial representation, in each mouse we correlated (Pearson’s r) the PV with the current neuronal activity for every time frame of the behavioral recording session. The guiding hypothesis was that correlations between neuronal activity and a hippocampal representation would be greater when the mouse was in the context from which the PV was constructed. To control for auto-correlations, we calculated PVs excluding the time bin with which the correlation was being calculated. In addition to dynamic PV correlations calculated by time frame, a mean population vector correlation was calculated for each mouse that considered the entire period (e.g. pre-shock or context A exposure).The difference between PV correlations was calculated to quantify the strength of one spatial PV over another (e.g. the difference of context A and context B spatial PVs during context A exposure). The guiding hypothesis was that the spatial PV of the presented context would have a greater correlation than the spatial PV of the non-presented context. These PV correlation differences were calculated either dynamically by time frame or during an entire context period.

To confirm significant PV correlation differences between groups with unequal cell numbers, we downsampled to match cell numbers between groups (**Figure S4**). This procedure was used in comparisons between vCA1 and dCA1, and conditioned and non-conditioned groups. For a given comparison, the group with more recorded cells was randomly downsampled to match the number of cells recorded in the comparison group with fewer cells. The rate maps and PVs were re-constructed using the downsampled cell population and the mean PV correlation was calculated for each mouse. This random downsampling cell selection and PV correlation analysis was performed 1000 times for each mouse. We then calculated the mean of the 1000 random iterations for each mouse and used that value to perform statistical analyses with the group that had fewer recorded cells.

#### Statistical analysis

Statistical analyses for behavioral and calcium imaging data were performed using custom MATLAB (Mathworks) scripts and Prism (GraphPad) software. Below is an inventory of parametric and non-parametric statistical tests used, for specific parameters and analyses refer to the figure legends. To compare behavior between conditioned and non-conditioned groups across time, a two-way repeated measures ANOVA adjusted for lack of sphericity by a Geisser-Greenhouse correction and with group and time as the factors was used. Changes in percent freezing during fear conditioning, discrimination test, and fear extinction were assessed with a one-way repeated measure ANOVA. All ANOVA tests with significant interactions were followed by a Bonferroni multiple comparison post hoc test. Statistical comparisons between two groups used a two-tailed Wilcoxon rank sum test, whereas within-group comparisons used either a Wilcoxon signed-rank test or a Kruskal Wallis test, followed by a Dunn multiple comparison test when there were greater than two conditions. Significant correlations were assessed using Pearson correlation. To assess the difference between two distributions a two-sample Kolmogorov-Smirnov test was used. To compare differences in the dynamics of context population vector correlations over time, pairwise comparisons between PV A and B correlation values were performed using Wilcoxon signed-rank tests. During fear extinction, the relationship of PV-A correlation during multiple context A exposures was assessed with the Friedman test. Throughout the behavioral tasks, the same mice were used on days 1 to 4. The only exception were two vCA1 non-conditioned mice that were excluded from day 1 due to calcium imaging complications that arose during the behavioral testing. All statistical tests used an alpha value of 0.05. Significance was determined as follows: * p<0.05, ** p<0.01, *** p<0.001, **** p<0.0001.

## Supplemental Information

**Figure S1 (related to Figure 1):**
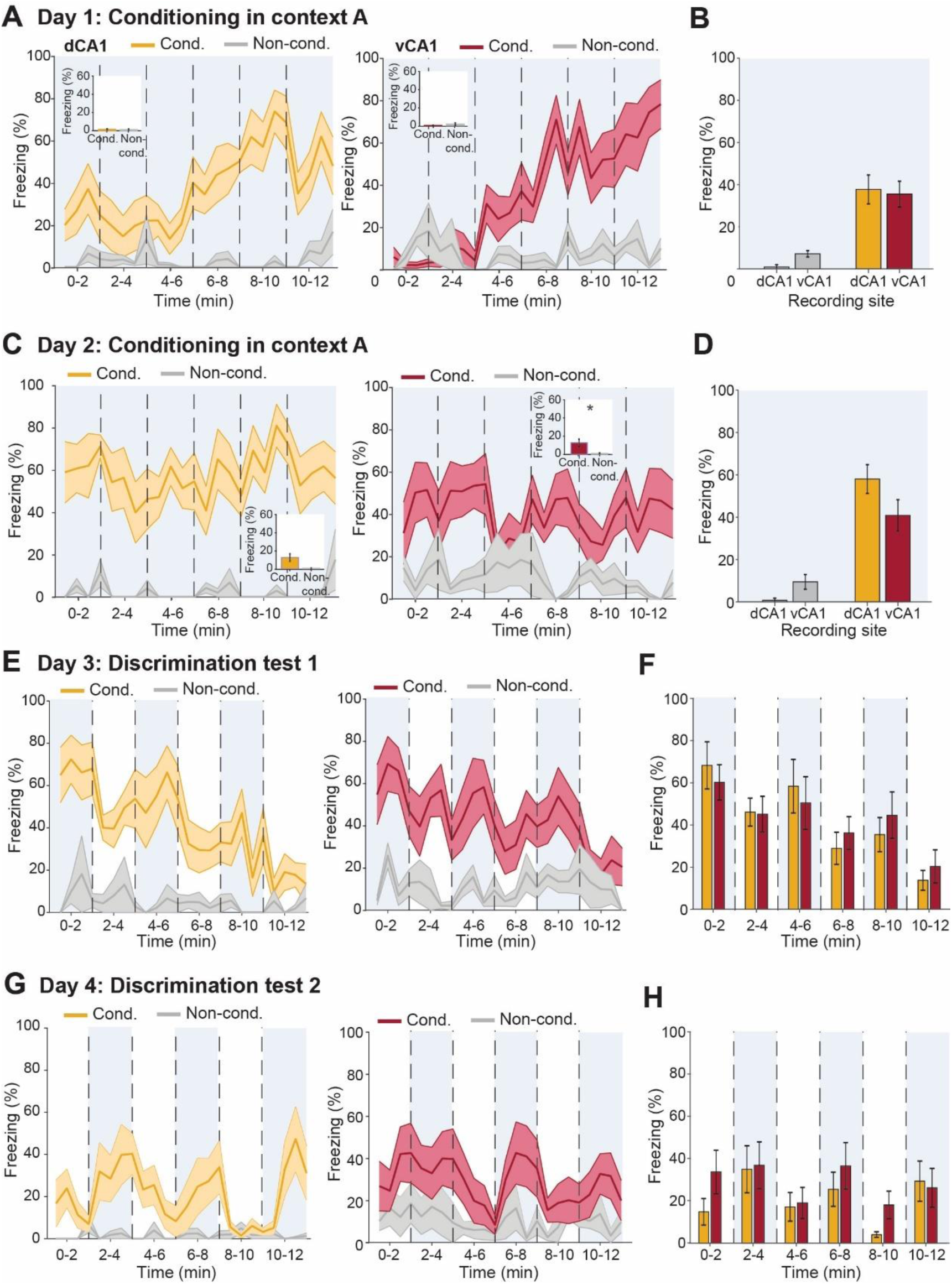
Freezing behavior during context fear discrimination task in dCA1 and vCA1 conditioned and non-conditioned groups. **(A)** Dynamics of freezing behavior on day 1 in dCA1 (left) and vCA1 (right) groups. Conditioned (Cond.) mice are shown in yellow for dCA1 (n = 8 mice) or in red for vCA1 (n = 8 mice). Non-conditioned (Non-cond.) mice are shown in gray (n = 4 mice for dCA1 and n = 6 for vCA1). Large panels correspond to context A (dCA1: two-way RM ANOVA: F_(1,10)_ = 16.33, p < 0.01; main effect of time F_(6.436, 64.36)_ = 0.688, p > 0.05; interaction F_(23, 230)_ = 1.765, p < 0.05 followed by a Bonferroni’s multiple comparison test; vCA1: two-way RM ANOVA: F_(1,12)_ = 16.46, p < 0.01; main effect of time F_(7.279, 87.35)_ = 2.138, p < 0.05; interaction F_(23, 276)_ = 2.056, p < 0.01 followed by a Bonferroni’s multiple comparison test). Panel inserts correspond to exposure in context B (dCA1: Wilcoxon rank-sum: rank = 53, p > 0.05; vCA1: Wilcoxon rank-sum: rank = 48, p > 0.05). **(B)** Freezing behavior of dCA1 and vCA1 conditioned and non-conditioned groups during day 1 (Wilcoxon rank-sum: rank = 13, p > 0.05) and conditioned mice (Wilcoxon rank-sum: rank = 64, p > 0.05). **(C)** Dynamics of freezing behavior on day 2 in dCA1 (left) and vCA1 (right) groups. Large panels correspond to context A (dCA1: two-way RM ANOVA: F_(1,10)_ = 52.33, p < 0.0001; main effect of time F_(5.99, 59.99)_ = 0.7619, p > 0.05; interaction F_(23, 230)_ = 0.4654, p > 0.05 followed by a Bonferroni’s multiple comparison test; vCA1: two-way RM ANOVA: F_(1,12)_ = 10.48, p < 0.001; main effect of time F_(8.55, 102.7)_ = 0.7296, p < 0.05; interaction F_(23, 276)_ = 0.9742, p > 0.05 followed by a Bonferroni’s multiple comparison test). Panel inserts correspond to exposure in context B (dCA1: Wilcoxon rank-sum: rank = 62, p > 0.05; vCA1: Wilcoxon rank-sum: rank = 73, p < 0.05). **(D)** Freezing behavior of dCA1 and vCA1 conditioned and non-conditioned groups during day 2 (Wilcoxon rank-sum: rank = 14, p > 0.05) and conditioned mice (Wilcoxon rank-sum: rank = 85, p > 0.05). **(E)** Dynamics of freezing behavior during discrimination test on day 3 in dCA1 (left, n = 8 conditioned mice, n = 4 non-conditioned mice, two-way RM ANOVA: F_(1,10)_ = 21.04, p < 0.001; main effect of time F_(6.301, 63.01)_ = 0.613, p > 0.05; interaction F_(23, 230)_ = 0.9453, p > 0.05 followed by a Bonferroni’s multiple comparison test) and vCA1 groups (right, n = 8 conditioned mice, n = 8 non-conditioned mice, two-way RM ANOVA: F_(1,14)_ = 15.79, p < 0.01; main effect of time F_(7.882, 110.4)_ = 1.003, p > 0.05; interaction F_(23, 322)_ = 1.047, p > 0.05 followed by a Bonferroni’s multiple comparison test). **(F)** Freezing behavior of dCA1 and vCA1 groups during discrimination test on day 3 (two-way RM ANOVA: F_(1,14)_ = 0.001, p > 0.05; main effect of time F_(3.623, 50.73)_ = 3.873, p < 0.01; interaction F_(5, 70)_ = 0.6134, p > 0.05 followed by a Bonferroni’s multiple comparison test). **(G)** Dynamics of freezing behavior during discrimination test on day 4 in dCA1 (left, two-way RM ANOVA: F_(1,10)_ = 4.101, p > 0.05; main effect of time F_(5.965, 59.65)_ = 0.6973, p > 0.05; interaction F_(23, 230)_ = 0.7808, p > 0.05 followed by a Bonferroni’s multiple comparison test) and vCA1 groups (right, two-way RM ANOVA: F_(1,14)_ = 3.069, p > 0.05; main effect of time F_(8.612, 120.6)_ = 1.033, p > 0.05; interaction F_(23, 322)_ = 0.9572, p > 0.05 followed by a Bonferroni’s multiple comparison test). **(H)** Freezing behavior comparison during discrimination test on Day 3 between dCA1 and vCA1 groups (two-way RM ANOVA: F_(1,14)_ = 0.66, p > 0.05; main effect of time F_(3.856, 53.99)_ = 2.167, p > 0.05; interaction F_(5, 70)_ = 0.7935, p > 0.05 following by a Bonferroni’s multiple comparison test). Data are expressed as the mean ± standard error of the mean.

**Figure S2 (related to Figure 2):**
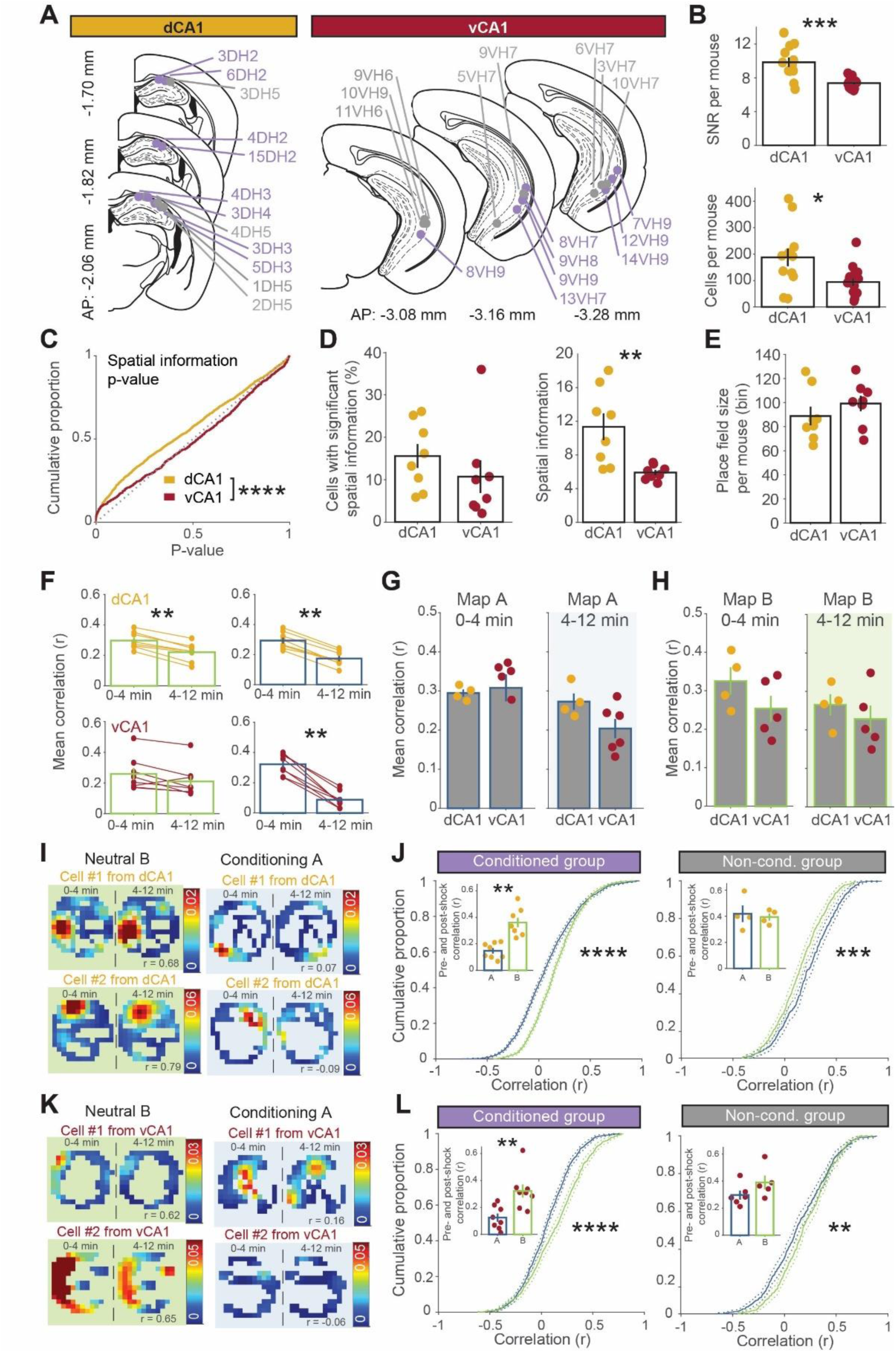
dCA1 encodes greater spatial information and vCA1 is most affected by conditioning. **(A)** Left, dCA1 GRIN lens placements for Ca2+ imaging of 8 experimental mice (purple dots indicate the bottom of the lens) and 4 control mice (in gray). Right, vCA1 GRIN lens placements of 8 experimental mice (in purple) and 8 control mice (in gray). **(B)** Top, average signal-to-noise ratio (SNR) per mouse is greater in dCA1 compared to vCA1 (Wilcoxon rank-sum: rank = 246, p < 0.001). Bottom, mean number of cells recorded per mouse is higher in dCA1 (Wilcoxon rank-sum: rank = 226.5, p < 0.05). **(C)** Cumulative distributions of spatial information p-values during context B exposure on day 1. dCA1 p-values were significantly left-shifted from the vCA1 distribution (Kolmogorov-Smirnov test, p < 0.0001, Kolmogorov-Smirnov stat = 0.11, n_dCA1_ = 1749 cells, n_vCA1_ = 627 cells) and both distributions were significantly different from the expected uniform non-spatially tuned distribution (diagonal dashed line) (dCA1: Kolmogorov-Smirnov test, p<0.0001, Kolmogorov-Smirnov stat = 0.12; vCA1: Kolmogorov-Smirnov test, p<0.05, Kolmogorov-Smirnov stat = 0.07). **(D)** Left, percent of neurons with significant spatial information p-values at a threshold of p = 0.05 in dCA1 and vCA1 are not significantly different (Wilcoxon rank-sum: rank = 83, p > 0.05). Right, mean spatial information values are significantly higher in dCA1 than in vCA1 (Wilcoxon rank-sum: rank = 95, p < 0.01). **(E)** Mean of place field sizes per mouse recorded in dCA1 and vCA1 during context B exposure on day 1 is not significantly different from each other (Wilcoxon rank-sum: rank = 59, p > 0.05). **(F)** Comparison of the mean values of the correlation with pre-shock PV calculated during pre-and post-shock periods of 0-4 min and 4-12 min for context B exposure. Top left, PV-B mean correlation in dCA1 was significantly reduced from minutes 0-4 to 4-12 (Wilcoxon signed-rank: rank = 36, p = 0.008). Top right, PV-A mean correlation in dCA1 was significantly reduced from minutes 0-4 to 4-12 (Wilcoxon signed-rank: rank = 36, p = 0.008). Bottom left, PV-B mean correlation in vCA1 was unchanged from minutes 0-4 to 4-12 (Wilcoxon signed-rank: rank = 31, p > 0.05). Bottom right, PV-A mean correlation in vCA1 was significantly reduced from minutes 0-4 to 4-12 (Wilcoxon signed-rank: rank = 36, p = 0.008). **(G)** Comparison of the mean values of the correlation with 0-4 min PV-A for dCA1 control mice (n = 4 mice) and vCA1 control mice (n = 5) during context A exposure without shock. Left, no significant difference in mean correlation values from 0-4 min in mice recorded from dCA1 and vCA1 (Wilcoxon rank-sum: rank = 17, p > 0.05). Right, no difference in mean correlation values of mice recorded in dCA1 and vCA1 from 4-12 min (Wilcoxon rank-sum: rank = 29, p > 0.05). **(H)** Comparison of the mean values of the correlation with 0-4 min PV-B for dCA1 control mice (n = 4 mice) and vCA1 control mice (n = 5) during context B exposure. Left, no significant difference in mean correlation values from 0-4 min in mice recorded from dCA1 and vCA1 (Wilcoxon rank-sum: rank = 26, p > 0.05). Right, no difference in mean correlation values of mice recorded in dCA1 and vCA1 from 4-12 min (Wilcoxon rank-sum: rank = 24, p > 0.05). **(I)** Examples of rate maps of cells recorded in dCA1 during context B and A on day 1. Within session rate map correlations between 0-4 and 4-12 min were performed for both contexts A (right) and B (left) to reflect pre-shock and post-shock exposure in context A. **(J)** Cumulative distributions of dCA1 place field map cross-correlations between pre-shock (or 0-4 min) and post-shock (4-12 min) during conditioning in context A (in blue) and context B exposure (in green). For dCA1 conditioned mice (n = 8), rate map correlations during conditioning in context A (in blue) were significantly left-shifted from rate map correlation distribution during context B exposure (in green) (Kolmogorov-Smirnov test, p<0.0001, Kolmogorov-Smirnov stat = 0.18, n_inA_ = 1859, n_inB_ = 1749). Left insert, mean of dCA1 rate map correlations between 0-4 min and 4-12 min with shock in context A (blue) and without shock in context B (green), indicated that the place fields were more stable without shock (Wilcoxon signed rank test: signed rank = 0, p < 0.01). Right, dCA1 non-conditioned mice (n = 4) had a lower difference in rate map correlation distributions (Kolmogorov-Smirnov test, p<0.001, Kolmogorov-Smirnov stat = 0.15, n_inA_ = 380, n_inB_ = 360), but there was no significant difference between the mean of rate map cross-correlations in context A (blue) and in context B (green), indicating that in the absence of shock place fields are stable (Wilcoxon signed rank test: signed rank = 5, p = 1). **(K)** Examples of rate maps as in **I** but recorded in vCA1. **(L)** Cumulative distributions of place field map cross-correlations as in **(J)** but recorded in vCA1. For vCA1 conditioned mice (n = 8), rate map correlations during conditioning in context A (in blue) were significantly left-shifted from rate map correlation distribution during context B exposure (in green) (Kolmogorov-Smirnov test, p<0.0001, Kolmogorov-Smirnov stat = 0.18, n_inA_ = 904, n_inB_ = 627). Left insert, mean of vCA1 rate map correlations between 0-4 min and 4-12 min with shock in context A (blue) and without shock in context B (green), indicated that the place fields were more stable without shock (Wilcoxon signed rank test: signed rank = 0, p < 0.01). Right, vCA1 non-conditioned mice (n = 5 or 6 for context B and A exposures respectively) had a lower difference in rate map correlation distributions (Kolmogorov-Smirnov test, p<0.01, Kolmogorov-Smirnov stat = 0.12, n_inA_ = 465, n_inB_ = 435), but there was no significant difference between the mean of rate map cross-correlations in context A (blue) and in context B (green) (Wilcoxon signed rank test: signed rank = 28, p = 0.18). Data are expressed as the mean ± standard error of the mean.

**Figure S3 (related to Figure 3):**
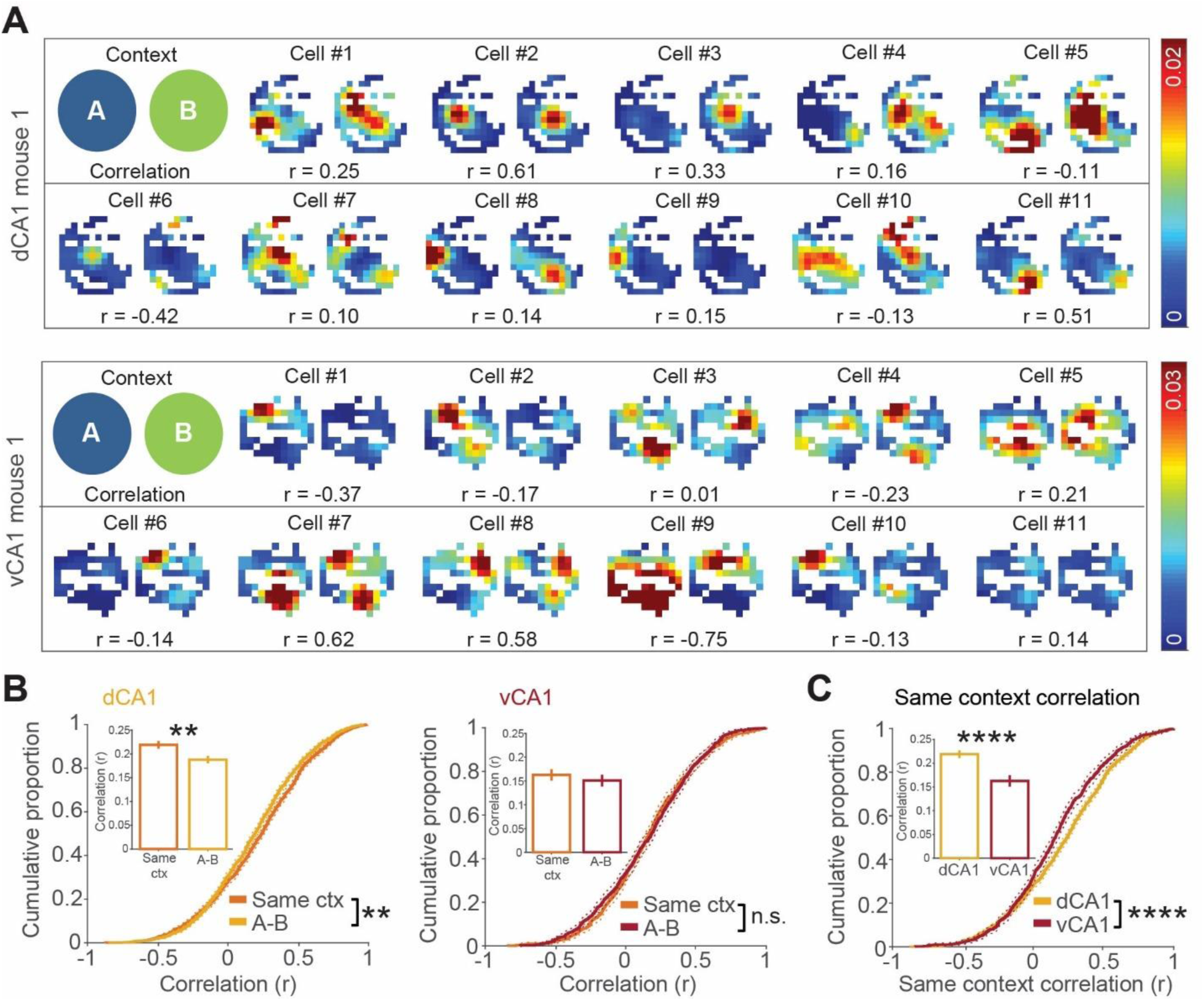
Remapping during context fear discrimination day 3. **(A)** Spatial rate maps were constructed by collapsing data across context A exposures during day 3; an identical method was used to construct context B rate maps. Spatial sampling was matched between contexts. Top, examples of dCA1 rate maps from context A (left) and context B (right). Bottom, examples of vCA1 rate maps from context A and context B. **(B)** Left, cumulative distributions of dCA1 place field map cross-correlations for same contexts (ctx) exposures (average of A-A and B-B place field correlations) (in orange) were significantly right-shifted relative to A-B correlations (in yellow) indicating that same context place fields were more stable (Kolmogorov-Smirnov test p = 0.0011, Kolmogorov-Smirnov stat = 0.0618, n = 1943 neurons). Left insert, mean dCA1 place field map cross-correlations of same context place field correlations and A-B correlations (Wilcoxon rank-sum: rank = 3677748, p = 0.0016). Right, cumulative distributions of vCA1 place field map cross-correlations were not significantly different between same context (orange) and A-B (red) exposures (Kolmogorov-Smirnov test p = 0.3835, Kolmogorov-Smirnov stat = 0.0456, n = 803 neurons). Right insert, mean vCA1 place field map cross-correlations of same context place field correlations and A-B correlations (Wilcoxon rank-sum: rank = 628934, p = 0.7727). **(C)** Comparison of the cumulative distributions for dCA1 or vCA1 place field map cross-correlations for same contexts exposures (Kolmogorov-Smirnov test p < 0.00011, Kolmogorov-Smirnov stat = 0.12, n_dCA1_ = 1943, n_vCA1_ = 803 neurons). Insert, mean place field map cross-correlations of same context place field correlations for dCA1 and vCA1, indicating that dCA1 place fields were more stable than vCA1 place fields (Wilcoxon rank-sum: rank = 2741728, p < 0.0001).

**Figure S4 (related to Figures 2, 3, 4):**
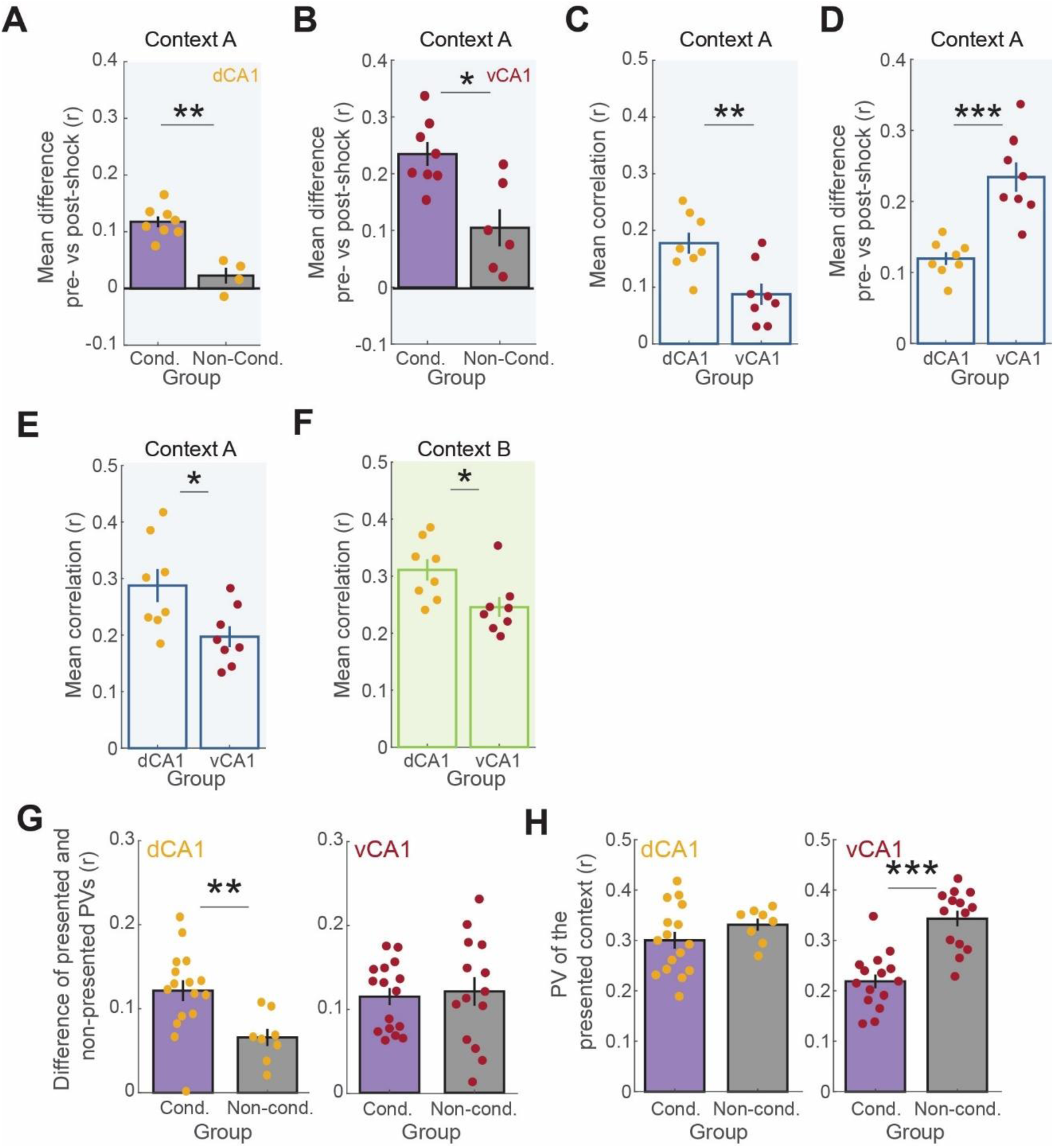
matching the number of neurons in population vector analyses between dCA1, vCA1, conditioned, and non-conditioned groups replicates region-specific and group-specific effects. **(A)** The number of neurons used in the dCA1 conditioned group (Cond.) was matched to the non-conditioned group (Non-cond.) and the post-shock PV was subtracted from pre-shock PV in context A on conditioning day 1 (see Figure 2E), replicating a significantly greater difference in the conditioned group (Wilcoxon rank-rum: rank = 68, p = 0.0040). **(B)** The number of neurons used in the vCA1 conditioned group was matched to the non-conditioned group and the post-shock PV was subtracted from pre-shock PV in context A on conditioning day 1 (see Figure 2G), replicating a significantly greater difference in the conditioned group (Wilcoxon rank-rum: rank = 79, p = 0.0127). **(C)** The number of neurons used in the dCA1 conditioned group was matched to the vCA1 conditioned group and the mean PV correlation for context A during the post-shock period was calculated on conditioning day 1 (see Figure 2H), replicating a significantly greater PV correlation in dCA1 (Wilcoxon rank-rum: rank = 92, p = 0.0104). **(D)** The number of neurons used in the dCA1 conditioned group was matched to the vCA1 conditioned group and the post-shock PV was subtracted from pre-shock PV in context A on conditioning day 1 (see Figure 2I), replicating a significantly greater difference in vCA1 conditioned group (Wilcoxon rank-rum: rank = 37, p = 0.0003). **(E)** The number of neurons used in the dCA1 conditioned group was matched to the vCA1 conditioned group and the mean PV correlation for context A during context A presentation was calculated on test day 3 (see Figure 3E), replicating a significantly greater PV correlation in dCA1 (Wilcoxon rank-rum: rank = 90, p = 0.0207). **(F)** The number of neurons used in the dCA1 conditioned group was matched to the vCA1 conditioned group and the mean PV correlation for context B during context B presentation was calculated on test day 3 (see Figure 3E), replicating a significantly greater PV correlation in dCA1 (Wilcoxon rank-rum: rank = 90, p = 0.0207). **(G)** Left, the number of neurons used in the dCA1 conditioned group was matched to the non-conditioned group and the difference of PV correlations for presented and non-presented contexts was calculated on test day 3 (see Figure 4E), replicating a significantly greater PV correlation in dCA1 conditioned group (Wilcoxon rank-rum: rank = 246, p = 0.0053). Right, the number of neurons used in the vCA1 conditioned group was matched to the non-conditioned group and the difference of PV correlations for presented and non-presented contexts was calculated on test day 3 (see Figure 4E), replicating a non-significant difference between vCA1 groups (Wilcoxon rank-rum: rank = 244, p > 0.05). **(H)** Left, the number of neurons used in the dCA1 conditioned group was matched to the non-conditioned group and the PV correlations for the presented contexts were calculated on test day 3 (see Figure 4F), replicating a non-significant difference between dCA1 groups (Wilcoxon rank- rum: rank = 180, p > 0.05). Right, the number of neurons used in the vCA1 conditioned group was matched to the non-conditioned group and the PV correlations for the presented contexts were calculated on test day 3 (see Figure 4F), replicating a significantly reduced correlation in vCA1 conditioned group (Wilcoxon rank-rum: rank = 148, p = 0.0004). Data are expressed as the mean ± standard error of the mean.

**Figure S5 (related to Figure 6):**
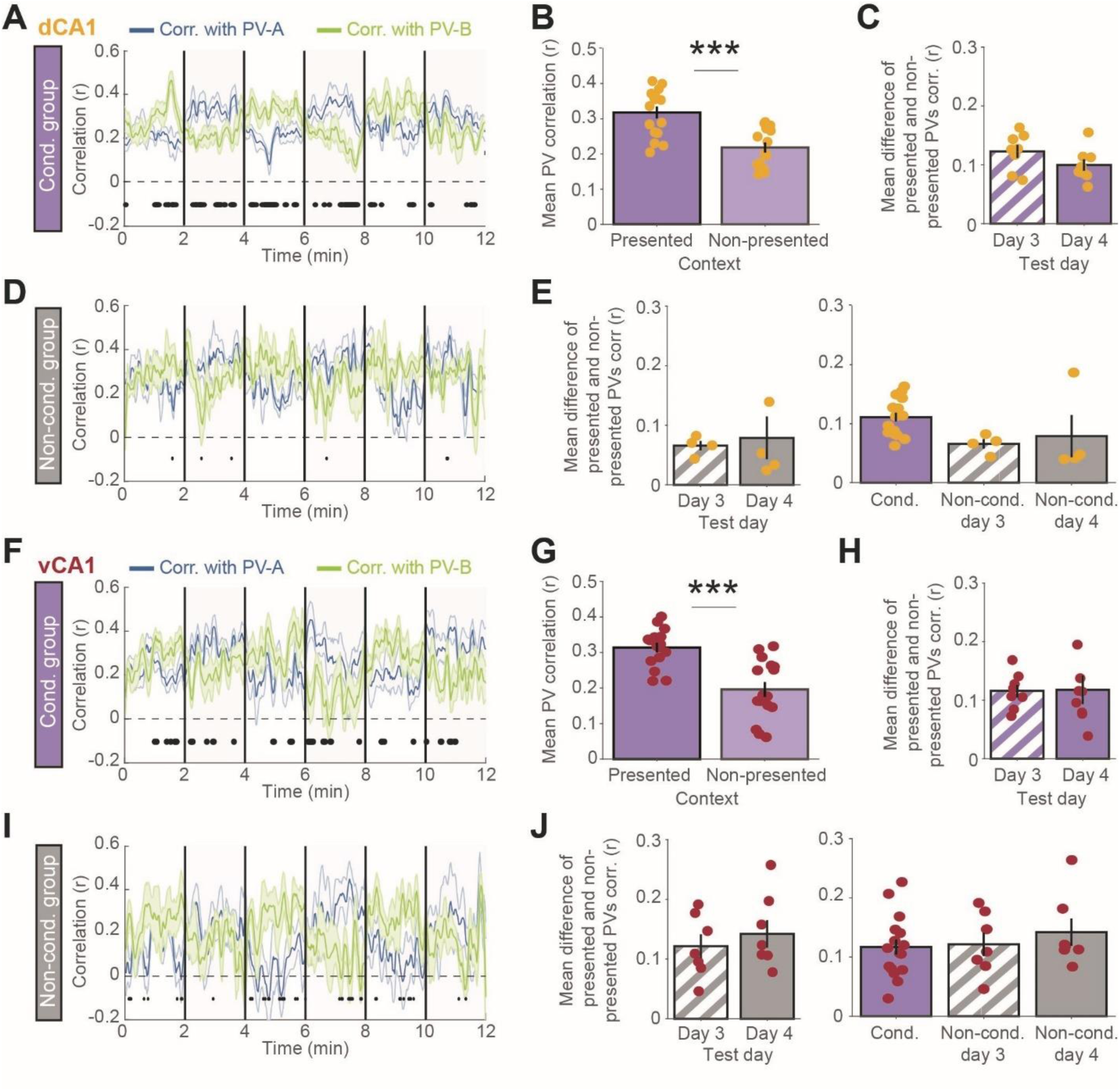
Discrimination test on day 4. **(A)** Dynamics of dCA1 population vector correlations with PV-A and PV-B in conditioned (Cond.) group (n = 8 mice) during day 4 discrimination test. Blue shaded regions indicate context A exposure, white region is context B. Black dots indicate timepoints of significant (Wilcoxon signed-rank test, p < 0.05) pairwise comparisons between PV-A and B correlation values. **(B)** dCA1 correlations with PV of the presented context (i.e. either PV-A during presentations of context A or PV-B during presentations of context B) were significantly greater than the PV correlations of the non-presented context (Wilcoxon signed rank test: signed rank = 136, p < 0.001). **(C)** Difference of the mean PV correlation of the presented context and the mean PV correlation of the non-presented context during days 3 and 4 in the dCA1 conditioned group (Wilcoxon rank test: rank = 30, p > 0.05). **(D)** Dynamics of population vector correlations with PV-A and PV-B as in **(A)** but recording in the dCA1 non-conditioned (Non-cond.) group (n = 4 mice). **(E)** Left, difference of the mean PV correlation of the presented context and the mean PV correlation of the non-presented context during days 3 and 4 in the dCA1 non-conditioned group (Wilcoxon rank test: rank = 5, p > 0.05). Right, comparison of the mean correlation difference between conditioned and non-conditioned groups in dCA1 (Kruskal-Wallis test: rank = 6.655, p = 0.04 followed by a Dunn’s multiple comparison test, post-test revealed no significant difference). **(F)** Dynamics of population vector correlations of PV-A and PV-B as in **(A)** but recording in the vCA1 conditioned group (n = 8 mice). **(G)** vCA1 correlations with PV of the presented context were significantly greater than the PV correlations of the non-presented context (Wilcoxon signed rank test: signed rank = 134, p < 0.001). **(H)** Difference of the mean PV correlation of the presented context and the mean PV correlation of the non-presented context during days 3 and 4 in the vCA1 conditioned group (Wilcoxon rank test: rank = 15, p > 0.05). **(I)** Dynamics of population vector correlations with PV-A and PV-B as in **(A)** but recording in the vCA1 non-conditioned group (n = 7 mice). **(J)** Left, difference of the mean PV correlation of the presented context and the mean PV correlation of the non-presented context during days 3 and 4 in the vCA1 non-conditioned group (Wilcoxon rank test: rank = 10, p > 0.05). Right, comparison of the mean correlation difference between conditioned and non-conditioned groups in vCA1 (Kruskal-Wallis test: rank = 1.024, p > 0.05). All data are expressed as the mean ± standard error of the mean.

**Figure S6 (related to Figure 6):**
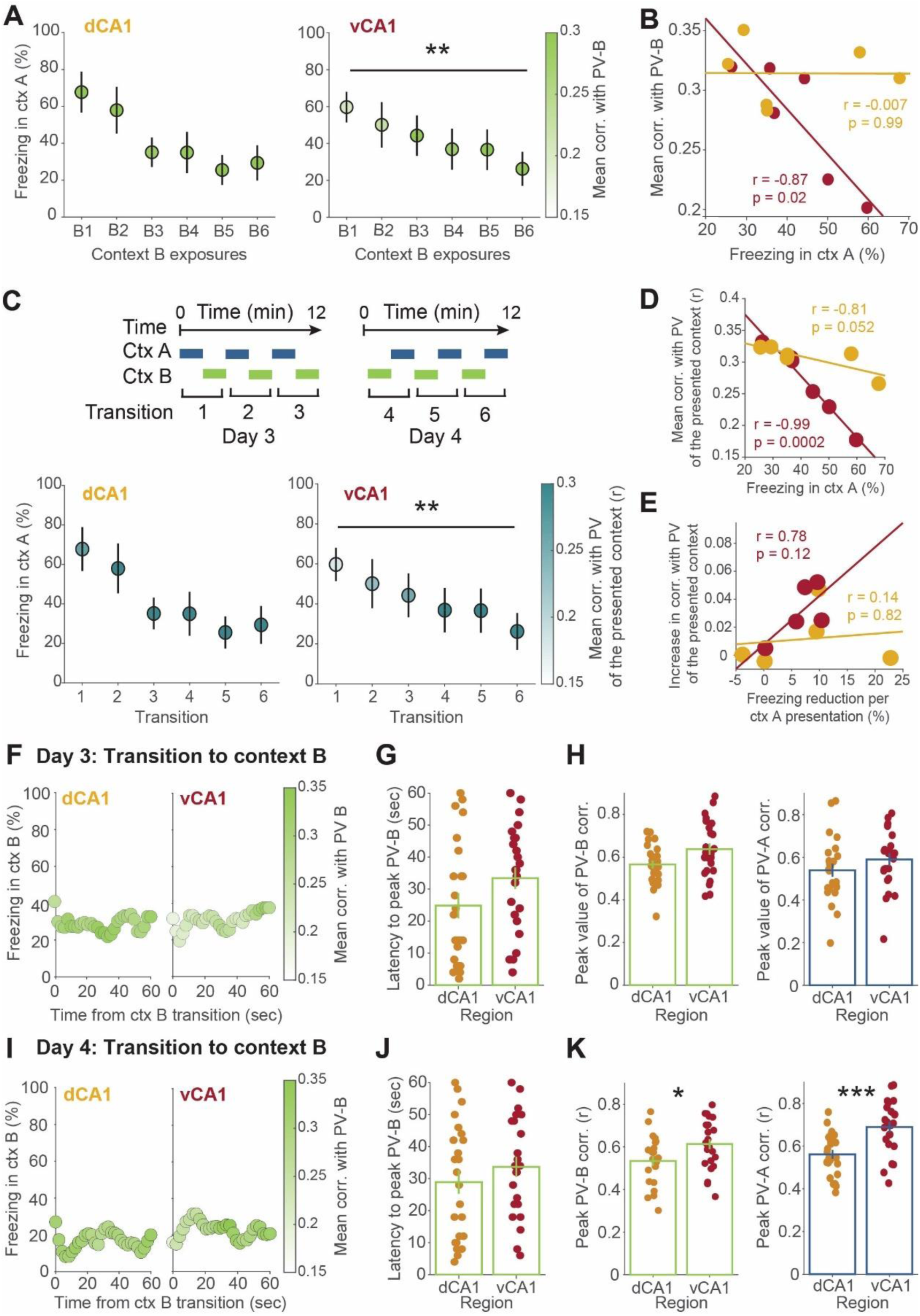
Fear extinction reverses conditioning-induced representational similarity in vCA1. **(A)** Mean correlation with PV-B during context (ctx) B exposure and freezing behavior in context A during discrimination tests on day 3 (B1-B2-B3) and 4 (B4-B5-B6) for dCA1 (left, n = 8 mice) and vCA1 (right, n = 8 mice). The mean correlation (corr.) with PV-B decreased during context B exposures in vCA1, but not in dCA1 (dCA1: Friedman’s test: F = 10.9, p = 0.0543; vCA1: Friedman’s test: F = 15.7, p < 0.01). **(B)** Correlation between the mean PV-B correlation values and freezing in context A (dCA1: correlation coefficient r = −0.007, p > 0.05, n = 6 context A exposures; vCA1: correlation coefficient r = −0.87, p < 0.05, n = 6 context A exposures). **(C)** Top, schematic of the transition numbers across test days 3 and 4. Bottom left, mean correlation of PV-A and PV-B during transitions into either context A or B and freezing behavior in context A during discrimination tests on days 3 and 4 for dCA1 (left, n = 8 mice) and vCA1 (bottom right, n = 8 mice). The mean correlation with PV-A and PV-B decreased over time in vCA1, but not in dCA1 (vCA1: Friedman’s test: F = 18.8, p < 0.01; dCA1: Friedman’s test: F = 6.07, p = 0.3). **(D)** Relationship between the mean correlation values of the PV-A and PV-B in **(C)** and freezing behavior in context A (dCA1: correlation coefficient r = −0.81, p > 0.05, n = 6 context transitions; vCA1: correlation coefficient r = −0.99, p < 0.001, n = 6 context transitions). **(E)** Correlation between the increase in PV A-B correlation and the decrease in freezing throughout context A transitions (dCA1: correlation coefficient r = 0.14, p > 0.05, n = 5 ITI; vCA1: correlation coefficient r = 0.78, p > 0.05, n = 5 ITI). **(F)** Dynamics of PV-B from the transition to context B during day 3 discrimination test for conditioned mice recorded in dCA1 or vCA1. **(G)** Latency to peak PV-B during transitions to context B on day 3 (n = 24, 8 mice for 3 context B transitions, Wilcoxon rank test: rank =503.5, p > 0.05). **(H)** Maximum peak values of PV-B (left) or PV-A (right) correlations during transition to context B or A respectively on day 3 (PV-B: Wilcoxon rank test: rank = 408, p = 0.065; PV-A: Wilcoxon rank test: rank = 508, p = 0.151). **(I)** Dynamics of PV-B from the transition to context B during day 4 discrimination test for conditioned mice recorded in dCA1 or vCA1. **(J)** Latency to peak PV-B during transitions to context B on day 4 (Wilcoxon rank test: rank =542.5, p > 0.05). **(K)** Maximum peak values of PV-B (left) or PV-A (right) correlations during transition to context B or A respectively on day 4 (PV B: Wilcoxon rank test: rank = 480, p < 0.05; PV A: Wilcoxon rank test: rank = 421, p < 0.001). All data are expressed as the mean ± standard error of the mean.

## Notes

### Competing Interest Statement

The authors have declared no competing interest.

## References

1. O’Keefe, J., and Burgess, N. (1996). Geometric determinants of the place fields of hippocampal neurons. Nature 381, 425–428. 10.1038/381425a0.

2. Muller, R.U., Kubie, J.L., and Ranck, J.B. (1987). Spatial firing patterns of hippocampal complex-spike cells in a fixed environment. J. Neurosci. 7, 1935–1950. 10.1523/JNEUROSCI.07-07-01935.1987.

3. Knierim, J.J., Kudrimoti, H.S., and McNaughton, B.L. (1995). Place cells, head direction cells, and the learning of landmark stability. J. Neurosci. 15, 1648–1659. 10.1523/JNEUROSCI.15-03-01648.1995.

4. Markus, E.J., Qin, Y.L., Leonard, B., Skaggs, W.E., McNaughton, B.L., and Barnes, C.A. (1995). Interactions between location and task affect the spatial and directional firing of hippocampal neurons. J. Neurosci. 15, 7079–7094. 10.1523/JNEUROSCI.15-11-07079.1995.

5. Wood, E.R., Dudchenko, P.A., Robitsek, R.J., and Eichenbaum, H. (2000). Hippocampal neurons encode information about different types of memory episodes occurring in the same location. Neuron 27, 623–633. 10.1016/s0896-6273(00)00071-4.

6. Shapiro, M.L., Tanila, H., and Eichenbaum, H. (1997). Cues that hippocampal place cells encode: Dynamic and hierarchical representation of local and distal stimuli. Hippocampus 7, 624–642. 10.1002/(SICI)1098-1063(1997)7:6<624::AID-HIPO5>3.0.CO;2-E.

7. Hollup, S.A., Molden, S., Donnett, J.G., Moser, M.-B., and Moser, E.I. (2001). Accumulation of Hippocampal Place Fields at the Goal Location in an Annular Watermaze Task. J. Neurosci. 21, 1635–1644. 10.1523/JNEUROSCI.21-05-01635.2001.

8. Bower, M.R., Euston, D.R., and McNaughton, B.L. (2005). Sequential-Context-Dependent Hippocampal Activity Is Not Necessary to Learn Sequences with Repeated Elements. J. Neurosci. 25, 1313–1323. 10.1523/JNEUROSCI.2901-04.2005.

9. Moita, M.A.P., Rosis, S., Zhou, Y., LeDoux, J.E., and Blair, H.T. (2004). Putting Fear in Its Place: Remapping of Hippocampal Place Cells during Fear Conditioning. J. Neurosci. 24, 7015–7023. 10.1523/JNEUROSCI.5492-03.2004.

10. Knierim, J.J., and Neunuebel, J.P. (2016). Tracking the flow of hippocampal computation: Pattern separation, pattern completion, and attractor dynamics. Neurobiology of Learning and Memory 129, 38–49. 10.1016/j.nlm.2015.10.008.

11. Stachenfeld, K.L., Botvinick, M.M., and Gershman, S.J. (2017). The hippocampus as a predictive map. Nat Neurosci 20, 1643–1653. 10.1038/nn.4650.

12. Buzsáki, G., and Moser, E.I. (2013). Memory, navigation and theta rhythm in the hippocampal-entorhinal system. Nature Neuroscience 16, 130–138. 10.1038/nn.3304.

13. Leutgeb, S., and Leutgeb, J.K. (2007). Pattern separation, pattern completion, and new neuronal codes within a continuous CA3 map. Learn Mem 14, 745–757. 10.1101/lm.703907.

14. Jezek, K., Henriksen, E.J., Treves, A., Moser, E.I., and Moser, M.-B. (2011). Theta-paced flickering between place-cell maps in the hippocampus. Nature 478, 246–249. 10.1038/nature10439.

15. Skaggs, W.E., and McNaughton, B.L. (1998). Spatial Firing Properties of Hippocampal CA1 Populations in an Environment Containing Two Visually Identical Regions. J. Neurosci. 18, 8455–8466. 10.1523/JNEUROSCI.18-20-08455.1998.

16. Anderson, M.I., and Jeffery, K.J. (2003). Heterogeneous modulation of place cell firing by changes in context. J Neurosci 23, 8827–8835.

17. Leutgeb, S., Leutgeb, J.K., Barnes, C.A., Moser, E.I., McNaughton, B.L., and Moser, M.-A. (2005). Independent codes for spatial and episodic memory in hippocampal neuronal ensembles. Science 309, 619–623. 10.1126/science.1114037.

18. Liu, X., Ramirez, S., Pang, P.T., Puryear, C.B., Govindarajan, A., Deisseroth, K., and Tonegawa, S. (2012). Optogenetic stimulation of a hippocampal engram activates fear memory recall. Nature 484, 381–385. 10.1038/nature11028.

19. Tayler, K.K., Tanaka, K.Z., Reijmers, L.G., and Wiltgen, B.J. (2013). Reactivation of Neural Ensembles during the Retrieval of Recent and Remote Memory. Current Biology 23, 99–106. 10.1016/j.cub.2012.11.019.

20. Tanaka, K.Z., Pevzner, A., Hamidi, A.B., Nakazawa, Y., Graham, J., and Wiltgen, B.J. (2014). Cortical representations are reinstated by the hippocampus during memory retrieval. Neuron 84, 347–354. 10.1016/j.neuron.2014.09.037.

21. Grella, S.L., Fortin, A.H., Ruesch, E., Bladon, J.H., Reynolds, L.F., Gross, A., Shpokayte, M., Cincotta, C., Zaki, Y., and Ramirez, S. (2022). Reactivating hippocampal-mediated memories during reconsolidation to disrupt fear. Nat Commun 13, 4733. 10.1038/s41467-022-32246-8.

22. Tomar, A., Polygalov, D., Chattarji, S., and McHugh, T.J. (2015). The dynamic impact of repeated stress on the hippocampal spatial map. Hippocampus 25, 38–50. 10.1002/hipo.22348.

23. Bouton, M.E., Maren, S., and McNally, G.P. (2021). BEHAVIORAL AND NEUROBIOLOGICAL MECHANISMS OF PAVLOVIAN AND INSTRUMENTAL EXTINCTION LEARNING. Physiol Rev 101, 611–681. 10.1152/physrev.00016.2020.

24. O’Keefe, J., and Dostrovsky, J. (1971). The hippocampus as a spatial map. Preliminary evidence from unit activity in the freely-moving rat. Brain Research 34, 171–175. 10.1016/0006-8993(71)90358-1.

25. O’Keefe, J., and Nadel, L. (1978). The Hippocampus as a Cognitive Map (Oxford: Clarendon Press).

26. Latuske, P., Kornienko, O., Kohler, L., and Allen, K. (2018). Hippocampal Remapping and Its Entorhinal Origin. Front Behav Neurosci 11. 10.3389/fnbeh.2017.00253.

27. Bostock, E., Muller, R.U., and Kubie, J.L. (1991). Experience-dependent modifications of hippocampal place cell firing. Hippocampus 1, 193–205. 10.1002/hipo.450010207.

28. Fyhn, M., Hafting, T., Treves, A., Moser, M.-B., and Moser, E.I. (2007). Hippocampal remapping and grid realignment in entorhinal cortex. Nature 446, 190–194. 10.1038/nature05601.

29. Keinath, A.T., Wang, M.E., Wann, E.G., Yuan, R.K., Dudman, J.T., and Muzzio, I.A. (2014). Precise Spatial Coding is Preserved Along the Longitudinal Hippocampal Axis. Hippocampus 24, 1533–1548. 10.1002/hipo.22333.

30. Wang, M.E., Wann, E.G., Yuan, R.K., Álvarez, M.M.R., Stead, S.M., and Muzzio, I.A. (2012). Long-Term Stabilization of Place Cell Remapping Produced by a Fearful Experience. J. Neurosci. 32, 15802–15814. 10.1523/JNEUROSCI.0480-12.2012.

31. Fanselow, M.S., and Dong, H.-W. (2010). Are the dorsal and ventral hippocampus functionally distinct structures? Neuron 65, 7–19. 10.1016/j.neuron.2009.11.031.

32. Royer, S., Sirota, A., Patel, J., and Buzsáki, G. (2010). Distinct Representations and Theta Dynamics in Dorsal and Ventral Hippocampus. J. Neurosci. 30, 1777–1787. 10.1523/JNEUROSCI.4681-09.2010.

33. Poucet, B., Thinus-Blanc, C., and Muller, R.U. (1994). Place cells in the ventral hippocampus of rats. NeuroReport 5, 2045.

34. Broadbent, N.J., and Clark, R.E. (2013). Remote context fear conditioning remains hippocampus-dependent irrespective of training protocol, training-surgery interval, lesion size, and lesion method. Neurobiol Learn Mem 106, 300–308. 10.1016/j.nlm.2013.08.008.

35. Ocampo, A.C., Squire, L.R., and Clark, R.E. (2017). Hippocampal area CA1 and remote memory in rats. Learn Mem 24, 563–568. 10.1101/lm.045781.117.

36. Sutherland, R.J., O’Brien, J., and Lehmann, H. (2008). Absence of systems consolidation of fear memories after dorsal, ventral, or complete hippocampal damage. Hippocampus 18, 710–718. 10.1002/hipo.20431.

37. Ciocchi, S., Passecker, J., Malagon-Vina, H., Mikus, N., and Klausberger, T. (2015). Selective information routing by ventral hippocampal CA1 projection neurons. Science 348, 560–563. 10.1126/science.aaa3245.

38. Jung, M., Wiener, S., and McNaughton, B. (1994). Comparison of spatial firing characteristics of units in dorsal and ventral hippocampus of the rat. J. Neurosci. 14, 7347– 7356. 10.1523/JNEUROSCI.14-12-07347.1994.

39. Moser, M.B., Moser, E.I., Forrest, E., Andersen, P., and Morris, R.G. (1995). Spatial learning with a minislab in the dorsal hippocampus. Proceedings of the National Academy of Sciences 92, 9697–9701. 10.1073/pnas.92.21.9697.

40. Nadel, L. (1968). Dorsal and ventral hippocampal lesions and behavior. Physiology & Behavior 3, 891–900. 10.1016/0031-9384(68)90174-1.

41. Felix-Ortiz, A.C., Beyeler, A., Seo, C., Leppla, C.A., Wildes, C.P., and Tye, K.M. (2013). BLA to vHPC Inputs Modulate Anxiety-Related Behaviors. Neuron 79, 658–664. 10.1016/j.neuron.2013.06.016.

42. Forro, T., Volitaki, E., Malagon-Vina, H., Klausberger, T., Nevian, T., and Ciocchi, S. (2022). Anxiety-related activity of ventral hippocampal interneurons. Prog Neurobiol 219, 102368. 10.1016/j.pneurobio.2022.102368.

43. Jimenez, J.C., Berry, J.E., Lim, S.C., Ong, S.K., Kheirbek, M.A., and Hen, R. (2020). Contextual fear memory retrieval by correlated ensembles of ventral CA1 neurons. Nature Communications 11, 3492. 10.1038/s41467-020-17270-w.

44. Padilla-Coreano, N., Canetta, S., Mikofsky, R.M., Alway, E., Passecker, J., Myroshnychenko, M.V., Garcia-Garcia, A.L., Warren, R., Teboul, E., Blackman, D.R., et al. (2019). Hippocampal-prefrontal theta transmission regulates avoidance behavior. Neuron 104, 601–610.e4. 10.1016/j.neuron.2019.08.006.

45. Çavdaroğlu, B., Toy, J., Schumacher, A., Carvalho, G., Patel, M., and Ito, R. (2020). Ventral hippocampus inactivation enhances the extinction of active avoidance responses in the presence of safety signals but leaves discrete trial operant active avoidance performance intact. Hippocampus 30, 913–925. 10.1002/hipo.23202.

46. Nguyen, R., Koukoutselos, K., Forro, T., and Ciocchi, S. (2023). Fear extinction relies on ventral hippocampal safety codes shaped by the amygdala. Science Advances 9, eadg4881. 10.1126/sciadv.adg4881.

47. Marek, R., Jin, J., Goode, T.D., Giustino, T.F., Wang, Q., Acca, G.M., Holehonnur, R., Ploski, J.E., Fitzgerald, P.J., Lynagh, T., et al. (2018). Hippocampus-driven feed-forward inhibition of the prefrontal cortex mediates relapse of extinguished fear. Nat Neurosci 21, 384–392. 10.1038/s41593-018-0073-9.

48. Bast, T., Zhang, W.N., and Feldon, J. (2001). The ventral hippocampus and fear conditioning in rats. Different anterograde amnesias of fear after tetrodotoxin inactivation and infusion of the GABA(A) agonist muscimol. Exp Brain Res 139, 39–52. 10.1007/s002210100746.

49. Rudy, J.W., and Matus-Amat, P. (2005). The Ventral Hippocampus Supports a Memory Representation of Context and Contextual Fear Conditioning: Implications for a Unitary Function of the Hippocampus. Behavioral Neuroscience 119, 154–163. 10.1037/0735-7044.119.1.154.

50. Pentkowski, N.S., Blanchard, D.C., Lever, C., Litvin, Y., and Blanchard, R.J. (2006). Effects of lesions to the dorsal and ventral hippocampus on defensive behaviors in rats. European Journal of Neuroscience 23, 2185–2196. 10.1111/j.1460-9568.2006.04754.x.

51. Adhikari, A., Topiwala, M.A., and Gordon, J.A. (2010). Synchronized activity between the ventral hippocampus and the medial prefrontal cortex during anxiety. Neuron 65, 257–269. 10.1016/j.neuron.2009.12.002.

52. Rozeske, R.R., Jercog, D., Karalis, N., Chaudun, F., Khoder, S., Girard, D., Winke, N., and Herry, C. (2018). Prefrontal-Periaqueductal Gray-Projecting Neurons Mediate Context Fear Discrimination. Neuron 97, 898–910.e6. 10.1016/j.neuron.2017.12.044.

53. Aharoni, D., Khakh, B.S., Silva, A.J., and Golshani, P. (2019). All the light that we can see: a new era in miniaturized microscopy. Nat Methods 16, 11–13. 10.1038/s41592-018-0266-x.

54. Carrillo-Reid, L., and Yuste, R. (2020). What Is a Neuronal Ensemble? Oxford Research Encyclopedia of Neuroscience. 10.1093/acrefore/9780190264086.013.298.

55. Keinath, A.T., Nieto-Posadas, A., Robinson, J.C., and Brandon, M.P. (2020). DG–CA3 circuitry mediates hippocampal representations of latent information. Nature Communications 11, 3026. 10.1038/s41467-020-16825-1.

56. Bannerman, D.M., Rawlins, J.N.P., McHugh, S.B., Deacon, R.M.J., Yee, B.K., Bast, T., Zhang, W.-N., Pothuizen, H.H.J., and Feldon, J. (2004). Regional dissociations within the hippocampus—memory and anxiety. Neuroscience & Biobehavioral Reviews 28, 273–283. 10.1016/j.neubiorev.2004.03.004.

57. Goosens, K.A. (2011). Hippocampal regulation of aversive memories. Current Opinion in Neurobiology 21, 460–466. 10.1016/j.conb.2011.04.003.

58. Cenquizca, L.A., and Swanson, L.W. (2007). Spatial organization of direct hippocampal field CA1 axonal projections to the rest of the cerebral cortex. Brain Res Rev 56, 1–26. 10.1016/j.brainresrev.2007.05.002.

59. Cembrowski, M.S., Wang, L., Sugino, K., Shields, B.C., and Spruston, N. (2016). Hipposeq: a comprehensive RNA-seq database of gene expression in hippocampal principal neurons. eLife 5, e14997. 10.7554/eLife.14997.

60. Dougherty, K.A., Islam, T., and Johnston, D. (2012). Intrinsic excitability of CA1 pyramidal neurones from the rat dorsal and ventral hippocampus. J Physiol 590, 5707–5722. 10.1113/jphysiol.2012.242693.

61. Malik, R., Dougherty, K.A., Parikh, K., Byrne, C., and Johnston, D. (2016). Mapping the electrophysiological and morphological properties of CA1 pyramidal neurons along the longitudinal hippocampal axis. Hippocampus 26, 341–361. 10.1002/hipo.22526.

62. Moser, M.-B., and Moser, E.I. (1998). Functional differentiation in the hippocampus. Hippocampus 8, 608–619. 10.1002/(SICI)1098-1063(1998)8:6<608::AID-HIPO3>3.0.CO;2-7.

63. Gray, J.A., and McNaughton, N. (1983). Comparison between the behavioural effects of septal and hippocampal lesions: A review. Neuroscience and Biobehavioral Reviews 7, 119–188. 10.1016/0149-7634(83)90014-3.

64. Ruediger, S., Spirig, D., Donato, F., and Caroni, P. (2012). Goal-oriented searching mediated by ventral hippocampus early in trial-and-error learning. Nat Neurosci 15, 1563– 1571. 10.1038/nn.3224.

65. McDonald, R.J., Balog, R.J., Lee, J.Q., Stuart, E.E., Carrels, B.B., and Hong, N.S. (2018). Rats with ventral hippocampal damage are impaired at various forms of learning including conditioned inhibition, spatial navigation, and discriminative fear conditioning to similar contexts. Behav Brain Res 351, 138–151. 10.1016/j.bbr.2018.06.003.

66. Wang, X., Zhang, C., Szábo, G., and Sun, Q.-Q. (2013). Distribution of CaMKIIα expression in the brain in vivo, studied by CaMKIIα-GFP mice. Brain Res 1518, 9–25. 10.1016/j.brainres.2013.04.042.

67. Jimenez, J.C., Su, K., Goldberg, A.R., Luna, V.M., Biane, J.S., Ordek, G., Zhou, P., Ong, S.K., Wright, M.A., Zweifel, L., et al. (2018). Anxiety Cells in a Hippocampal-Hypothalamic Circuit. Neuron 97, 670–683.e6. 10.1016/j.neuron.2018.01.016.

68. Adhikari, A., Topiwala, M.A., and Gordon, J.A. (2011). Single units in the medial prefrontal cortex with anxiety-related firing patterns are preferentially influenced by ventral hippocampal activity. Neuron 71, 898–910. 10.1016/j.neuron.2011.07.027.

69. Herry, C., Ciocchi, S., Senn, V., Demmou, L., Müller, C., and Lüthi, A. (2008). Switching on and off fear by distinct neuronal circuits. Nature 454, 600–606. 10.1038/nature07166.

70. Oler, J.A., Penley, S.C., Sava, S., and Markus, E.J. (2008). Does the dorsal hippocampus process navigational routes or behavioral context? A single-unit analysis. European Journal of Neuroscience 28, 802–812. 10.1111/j.1460-9568.2008.06375.x.

71. Kim, E.J., Park, M., Kong, M.-S., Park, S.G., Cho, J., and Kim, J.J. (2015). Alterations of Hippocampal Place Cells in Foraging Rats Facing a “Predatory” Threat. Current Biology 25, 1362–1367. 10.1016/j.cub.2015.03.048.

72. Jin, S.-W., and Lee, I. (2021). Differential encoding of place value between the dorsal and intermediate hippocampus. Current Biology 31, 3053–3072.e5. 10.1016/j.cub.2021.04.073.

73. Farzanfar, D., Spiers, H.J., Moscovitch, M., and Rosenbaum, R.S. (2023). From cognitive maps to spatial schemas. Nat Rev Neurosci 24, 63–79. 10.1038/s41583-022-00655-9.

74. Kjelstrup, K.B., Solstad, T., Brun, V.H., Hafting, T., Leutgeb, S., Witter, M.P., Moser, E.I., and Moser, M.-B. (2008). Finite Scale of Spatial Representation in the Hippocampus. Science 321, 140–143. 10.1126/science.1157086.

75. Hopfield, J.J. (1982). Neural networks and physical systems with emergent collective computational abilities. Proceedings of the National Academy of Sciences 79, 2554–2558. 10.1073/pnas.79.8.2554.

76. Johnson, A., and Redish, A.D. (2007). Neural Ensembles in CA3 Transiently Encode Paths Forward of the Animal at a Decision Point. J. Neurosci. 27, 12176–12189. 10.1523/JNEUROSCI.3761-07.2007.

77. Kay, K., Chung, J.E., Sosa, M., Schor, J.S., Karlsson, M.P., Larkin, M.C., Liu, D.F., and Frank, L.M. (2020). Constant Sub-second Cycling between Representations of Possible Futures in the Hippocampus. Cell 180, 552–567.e25. 10.1016/j.cell.2020.01.014.

78. Tronson, N.C., Schrick, C., Guzman, Y.F., Huh, K.H., Srivastava, D.P., Penzes, P., Guedea, A.L., Gao, C., and Radulovic, J. (2009). Segregated Populations of Hippocampal Principal CA1 Neurons Mediating Conditioning and Extinction of Contextual Fear. J. Neurosci. 29, 3387–3394. 10.1523/JNEUROSCI.5619-08.2009.

79. Lacagnina, A.F., Brockway, E.T., Crovetti, C.R., Shue, F., McCarty, M.J., Sattler, K.P., Lim, S.C., Santos, S.L., Denny, C.A., and Drew, M.R. (2019). Distinct hippocampal engrams control extinction and relapse of fear memory. Nat Neurosci 22, 753–761. 10.1038/s41593-019-0361-z.

80. Mount, R.A., Sridhar, S., Hansen, K.R., Mohammed, A.I., Abdulkerim, M., Kessel, R., Nazer, B., Gritton, H.J., and Han, X. (2021). Distinct neuronal populations contribute to trace conditioning and extinction learning in the hippocampal CA1. eLife 10, e56491. 10.7554/eLife.56491.

81. Lee, C., Lee, B.H., Jung, H., Lee, C., Sung, Y., Kim, H., Kim, J., Shim, J.Y., Kim, J., Choi, D.I., et al. (2023). Hippocampal engram networks for fear memory recruit new synapses and modify pre-existing synapses in vivo. Current Biology 33, 507–516.e3. 10.1016/j.cub.2022.12.038.

82. Wang, M.E., Yuan, R.K., Keinath, A.T., Álvarez, M.M.R., and Muzzio, I.A. (2015). Extinction of Learned Fear Induces Hippocampal Place Cell Remapping. J. Neurosci. 35, 9122–9136. 10.1523/JNEUROSCI.4477-14.2015.

83. Lopes, G., Bonacchi, N., Frazão, J., Neto, J.P., Atallah, B.V., Soares, S., Moreira, L., Matias, S., Itskov, P.M., Correia, P.A., et al. (2015). Bonsai: an event-based framework for processing and controlling data streams. Front. Neuroinform. 9. 10.3389/fninf.2015.00007.

84. Pnevmatikakis, E.A., and Giovannucci, A. (2017). NoRMCorre: An online algorithm for piecewise rigid motion correction of calcium imaging data. Journal of Neuroscience Methods 291, 83–94. 10.1016/j.jneumeth.2017.07.031.

85. Pnevmatikakis, E.A., Soudry, D., Gao, Y., Machado, T.A., Merel, J., Pfau, D., Reardon, T., Mu, Y., Lacefield, C., Yang, W., et al. (2016). Simultaneous Denoising, Deconvolution, and Demixing of Calcium Imaging Data. Neuron 89, 285–299. 10.1016/j.neuron.2015.11.037.

86. Zhou, P., Resendez, S.L., Rodriguez-Romaguera, J., Jimenez, J.C., Neufeld, S.Q., Giovannucci, A., Friedrich, J., Pnevmatikakis, E.A., Stuber, G.D., Hen, R., et al. (2018). Efficient and accurate extraction of in vivo calcium signals from microendoscopic video data. eLife 7, e28728. 10.7554/eLife.28728.

87. Friedrich, J., Zhou, P., and Paninski, L. (2017). Fast online deconvolution of calcium imaging data. PLOS Computational Biology 13, e1005423. 10.1371/journal.pcbi.1005423.

88. Giovannucci, A., Friedrich, J., Gunn, P., Kalfon, J., Brown, B.L., Koay, S.A., Taxidis, J., Najafi, F., Gauthier, J.L., Zhou, P., et al. (2019). CaImAn an open source tool for scalable calcium imaging data analysis. eLife 8, e38173. 10.7554/eLife.38173.

89. Sík, A., Hájos, N., Gulácsi, A., Mody, I., and Freund, T.F. (1998). The absence of a major Ca2+ signaling pathway in GABAergic neurons of the hippocampus. Proceedings of the National Academy of Sciences 95, 3245–3250. 10.1073/pnas.95.6.3245.

90. Muller, R. (1996). A Quarter of a Century of Place Cells. Neuron 17, 813–822. 10.1016/S0896-6273(00)80214-7.

91. Epsztein, J., Brecht, M., and Lee, A.K. (2011). Intracellular Determinants of Hippocampal CA1 Place and Silent Cell Activity in a Novel Environment. Neuron 70, 109– 120. 10.1016/j.neuron.2011.03.006.

92. Skaggs, W., McNaughton, B., and Gothard, K. (1992). An Information-Theoretic Approach to Deciphering the Hippocampal Code. In Advances in Neural Information Processing Systems (Morgan-Kaufmann).

